# Contact-based transfer of thymidylate promotes collective tumor growth

**DOI:** 10.1101/2024.07.31.606056

**Authors:** Mohammad A. Siddiqui, Anne Mette A. Rømer, Luisa Pinna, Vignesh Ramesh, Sarah S. Møller, Paradesi N. Gollavilli, Adriana M. Turtos, Steffi Varghese, Lærke B. Guldborg, Nagendra P. Palani, Francesca Napoli, Marco Volante, Luis Córdova Bahena, Marco Velasco, Kumar Somyajit, Pawel Swietach, Paolo Ceppi

## Abstract

Sustained cell proliferation is a fundamental hallmark of cancer, yet its mechanism remains elusive, particularly in context of heterogenous tumor organization, intercellular interactions, and metabolite exchanges. In this study, we uncover a mechanism of tumor growth via *collective proliferation*, where cells connected by gap junctions enable equilibration of thymidylate (dTMP) to allow the proliferation of cells lacking canonical *de novo* dTMP biosynthesis and salvage driven by thymidylate synthase (*TYMS*) and thymidine kinase 1 (*TK1*) enzymes, respectively. Collective proliferation is driven by dTMP-proficient cancer cells alongside cells of varied origins, such as macrophages and endothelial cells. The mechanism is also observed in clinical samples and is validated in a genetic mouse model of lung cancer harboring dual *Tyms*/*Tk1* tumor-specific knockout, in which tumors grow despite lacking enzymatic dTMP synthesis and tumor progression is opposed by gap junction inhibition. Data further hint that this mechanism is critical driver of tumor pathophysiology, influencing key processes such as senescence, genomic instability and drug resistance. These findings revise the current dogma of ubiquitous nucleotide biosynthesis in each proliferating cancer cell in a tumor and suggest that a programmed dTMP distribution maintains collective tumor growth. This mechanism could be exploited in cancer therapy.

## Introduction

Sustained cell proliferation, a key cancer hallmark, relies on a continuous supply of nucleotides either through multistep *de novo* biosynthesis or salvage of free nucleosides^1^. The current dogma of tumor growth emphasizes the universal activation of pyrimidine synthesis^2^, thereby propagating an energy-exhaustive growth model. However, tumors are genetically heterogeneous entities^3^ and cells with undetected levels of critical pyrimidine genes are frequently reported in clinical samples^4,5^. The contribution and significance of these cells in cancer growth has never been identified.

Recent studies have suggested that intercellular genetic heterogeneity, caused by disruption in key metabolic pathway genes, can be compensated by the equilibration of metabolites between electrically coupled cells forming a syncytium^6^. To investigate if pyrimidines could be shared in a similar way, we investigated proliferation dynamics between cells with active expression of nucleotide metabolism genes and cells with CRISPR/Cas9-mediated ablation of these genes. Strikingly, our findings reveal a compensatory mechanism where cells undergo collective proliferation, mediated by sharing of thymidylate among cells. These results, supported by *in vivo* models and clinical data, challenge the essentiality of intrinsically-driven TS/TK1-dependent proliferation, introducing a new paradigm in tumor growth with significant therapeutic implications.

### Intercellular dTMP transfer supports cancer cell proliferation

To maintain a continuous dTMP supply for DNA replication and proliferation, cancer cells primarily convert deoxyuridine monophosphate (dUMP) to deoxythymidine monophosphate (dTMP, thymidylate) *de novo* by thymidylate synthase (TS, gene name *TYMS*) or salvage extracellular thymidine using thymidine kinase-1 (TK1) (**Fig. 1A**). We used CRISPR/Cas9 to ablate *TYMS* or *TK1* in non-small cell lung cancer (NSCLC) cell lines. TS-KO reduced thymidylate levels, as quantified by mass-spectroscopy (**Extended Fig. 1A-B**), and induced absolute growth arrest but was not lethal. Compared to the parental cells, TS-KO cells were enlarged (**Extended Fig. 1C**) and proliferated only with supplementation of dTMP, but not dUMP **(Extended Fig. 1D)**, an effect observed with different non-overlapping gRNA sequences (**Extended Fig. 1E-F**). Furthermore, when testing different combinations of deoxyribonucleotides, only a combination containing dTTP (deoxythymidine triphosphates) re-established proliferation in TS-KO cells (**Extended Fig. 1G**). On contrary, knockout of *TK1* minimally affected cell growth (**Extended Fig. 2A-B**), implying that proliferation *in vitro* is primarily supported by TS-dependent *de novo* dTMP synthesis. Thereby, TS-KO cells represent a growth-deficient cell model that is entirely dependent on exogenous dTMP for proliferation.

**Figure 1.**
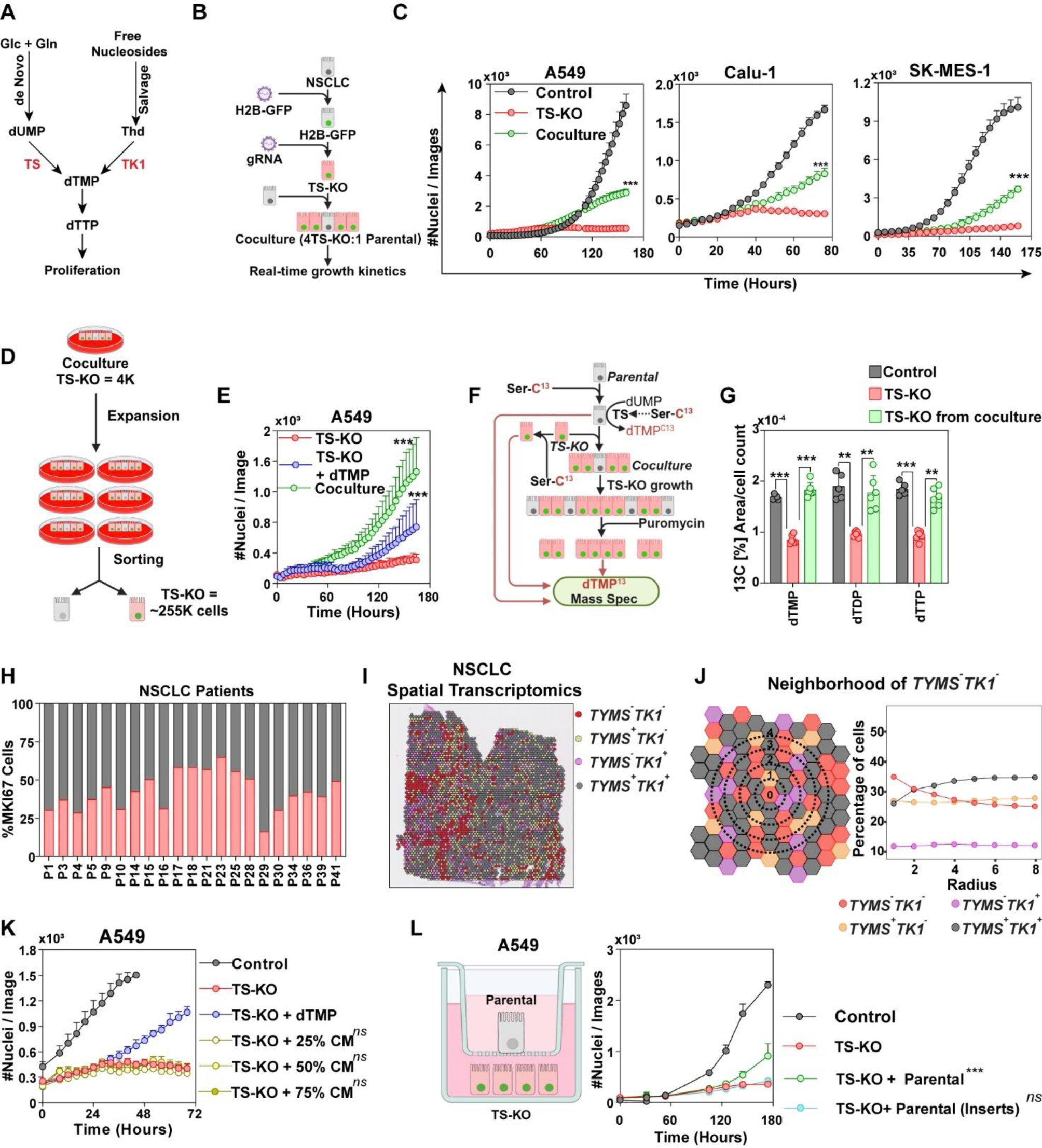
Thymidylate (dTMP) sharing in NSCLC cells. **(A)** Cancer cells synthesize dTMP either de novo by assimilating glucose and glutamine and converting dUMP to dTMP by TS mediated enzymatic conversion, or by TK-1 mediated salvaging of free thymidine (Thd). **(B)** NSCLC cell lines (A549, Calu-1 and SK-MES-1) expressing nuclear H2B-GFP, were knocked out for *TYMS* by CRISPR/Cas9 and cocultured with their respective parental cells (expressing TS) in a ratio of 4:1. Proliferation of GFP^+^ TS-KO cells was monitored using Incucyte S3. **(C)** The growth of TS-KO cells in coculture with TS-expressing parental. As experimental controls, GFP-tagged cells were transduced with vector containing a non-targeting sequence. The data are mean ± SD of three replicates and the statistical test is a Two-Way ANOVA test with Tukey’s multiple comparison test correction. **(D)** A549 TS-KO cells were cocultured with parental cells for 10 passages and sorted based on GFP expression. Cell counts at initial and final passage are indicated. **(E)** Sorted cells from (D) were seeded in a 96-well plate and their dependency on exogenous dTMP was quantified. The data are mean ± SD of three replicates and statistical test is Two-Way ANOVA test with Tukey’s multiple comparison test correction. **(F)** A549 cells were cocultured in DMEM supplemented with Serine-C^13^ for 24 hours to enrich the cells for dTMP^C13^ (A549^C13^). Puromycin resistant A549 TS-KO cells were cocultured with A549^C13^ parental cells and selected with puromycin after their growth. dTMP^C13^, dTDP^C13^ and, dTTP^C13^ were quantified in selected TS-KO cells. A549^C13^ and TS-KO cells cultured in Ser^C13^ for 24 hours were used as controls. See also the Extended Fig. 3. **(G)** Quantification of dTMP, dTDP, and dTTP tagged with C^13^ in the cells from (F). Bars represent mean ± SD of six replicates and statistical significance is calculated by One-Way ANOVA with Sidak’s multiple comparison. **(H)** Single-cell transcriptomic data from NSCLC patients showing contribution of cells lacking expression of *TYMS* and *TK1* (*TYMS*^-^*TK1*^-^) to the growth of tumors. *MKI67^+^* proliferative cells in *TYMS*^-^*TK1*^-^ population were compared to other populations (*TYMS*^+^*TK1*^-^, *TYMS*^-^*TK1*^+^ and *TYMS*^+^*TK1*^+^) for 21 patients (with at least 50 proliferative cells). Each bar represents a single patient. See also Extended Fig. 4. **(I)** Spatial transcriptomics data from a NSCLC biopsy, showing dispersion of *TYMS*^-^*TK1*^-^ cells with population with other genotypes. *TYMS^+^TK1^+^* = 1763 cells, *TYMS^+^TK1^-^* = 1000 cells, *TYMS^-^ TK1^+^* = 438 cells, *TYMS^-^TK1^-^* = 657 cells. **(J)** Neighborhood analysis of spatial transcriptomics data from (G). Taking *TYMS*^-^*TK1^-^* cells as center, percentages of different cell populations were calculated in successive radii. Percentages at each radius for each *TYMS*^-^*TK1^-^* cell was averaged and represented as a dot, representing tendency of a cell population to be located around these cells. **(K)** Proliferation kinetics of A549 TS-KO cells showing their inability to proliferate with supplementation of condition medium from A549 parental cells cultured for 48 hours. Each point represents mean ± SD of three replicates, statistical test is Two-Way ANOVA with Tukey’s multiple comparison. **(L)** Lack of growth of TS-KO cells in medium that is shared with the parental donor cells that were seeded on a transwell chamber, physically separating them from the TS-KO cells. Each point represents mean ± SD of three replicates, statistical test is Two-Way ANOVA with Tukey’s multiple comparison.

To monitor the proliferation dynamics of dTMP-deficient cells, *TYMS* was knocked out in NSCLC cell lines that express nuclear H2B-GFP, allowing for real-time tracking of GFP^+^ TS-KO cells in coculture with unlabeled parental cells (**Fig. 1B**). Unexpectedly, the TS-KO cells displayed restored growth when cocultured with parental cells, whereas TS-KO cells cultured alone failed to proliferate (**Fig. 1C)** and the morphology of cocultured TS-KO cells reverted to normal **(Extended Fig. 2C)**. The findings were confirmed using TS-KO cells generated by two other non-overlapping gRNAs in two different cell lines, where parental cell-dependent growth of TS-KO cells was replicated **(Extended Fig. 2D**). The quantification of the proliferative capacity of TS-KO cells was however limited by the spatial constraint from proliferative parental cells, and therefore a long-term coculture was established. A coculture of GFP^+^ TS-KO A549 cells and parental cells was expanded for 10 passages in sub-confluent conditions and the GFP^+^ TS-KO cells were flow-sorted and counted (**Fig. 1D**). When isolated, the sorted TS-KO cells failed to proliferate in absence of dTMP supplementation (**Fig. 1E**), demonstrating the stable nature of the dependency of TS-KO cells on parental cells for proliferation.

Next, to determine whether a similar observation could be made upon deficiency of other dNTPs, *RRM1* (to deplete all dNTPs) and *DTYMK* (to deplete pyrimidines) were knocked out in A549 cells (**Extended Fig. 2E-F)**. The growth defect arising from nucleotide depletion was, however, not restored in the coculture (**Extended Fig. 2G**), indicating the specificity of dTMP as the shared nucleotide between cells. To further demonstrate that dTMP equilibration across the coculture causes the proliferation of TS-KO cells, parental A549 cells were cultured in L-Serine-3-^13^C (Ser^13C^) because Serine’s C-3 is transferred to dUMP to synthesize dTMP (**Extended Fig. 3A**). Serine by itself did not reinstate the growth of TS-KO cells (**Extended Fig. 3B**). We also tested if nucleotides released by dead parental cells could support TS-KO cells proliferation but did not observe a rescue. (**Extended Fig. 3C-D**). Therefore, parental cells preincubated with Ser^13^ were cocultured with puromycin-resistant TS-KO cells. Once the re-growth of TS-KO cells was detected in coculture, parental cells were eliminated by puromycin treatment and dNTP^13C^ levels were quantified in proliferating TS-KO cells by mass spectrometry (**Fig. 1F**). As a result, dTMP levels were found re-equilibrated to the levels measured in parental cells, along with dTDP and dTTP (thymidine diphosphate and triphosphate respectively) (**Fig. 1G**), indicating that transfer of dTMP from the parental cells can rescue the growth defect caused by perturbation of the *de novo* dTMP synthesis.

To investigate if a collective modality of proliferation for dTMP-low cells exists also in tumors, single cell sequencing data of 40 NSCLC patients were pooled^7^ and the gene expression profiles of the various cell types were analyzed (**Extended Fig. 4A**). Interestingly, a significant proportion of the cancer cells lacked expression of *TYMS* (encoding TS) and *TK1* (*TYMS*^-^*TK1*^-^) in the cancer clusters (**Extended Fig. 4B**). In some patients *TYMS*^-^*TK1*^-^ comprised of more than 80% of the total cancer cell population (**Extended Fig. 4C**). Notably, when examining cells positive for the proliferation marker *MKI67* (in samples with over 50 proliferative cells) a substantial proportion of *TYMS^-^TK1^-^* cells was observed in all samples, reaching 50% of the total *MKI67*^+^ cells in certain cases (**Fig. 1H**). This highlights the significant contribution of *TYMS*^-^*TK1*^-^ cancer cells to tumor growth. Furthermore, *TYMS*^-^*TK1*^-^ cells frequently expressed other proliferative genes along with basic metabolic genes (**Extended Fig. 4D**), further validating the non-quiescent nature of this subpopulation. Next, a spatial transcriptomic dataset was analyzed to evaluate the distribution of *TYMS*^-^*TK1*^-^ cells in NSCLC tumors. The cells were found to be interspersed with other populations (*TYMS*^+^*TK1*^-^, *TYMS*^-^*TK1*^+^, *TYMS*^+^*TK1^+^*) (**Fig. 1I** and **Extended Fig. 5A**). Interestingly, a neighborhood analysis performed by calculating the proportion of different cell populations in successive radii revealed a remarkable tendency of *TYMS*^-^*TK1*^-^ cells to cluster near *TYMS*^+^*TK1*^+^ cells (**Fig. 1J**). Meanwhile, these observations were not found in the other groups of *TYMS*^-^*TK1*^+^ and *TYMS*^+^*TK1*^-^ populations, which appeared randomly located. *TYMS*^+^*TK1*^+^ cells showed the opposite trend (**Extended Fig. 5B**). This suggested that *TYMS*^-^*TK1*^-^ cells prefer easy access to the dTMP, potentially released into the extracellular space by the *TYMS*^+^*TK1*^+^ cells. To test this, we assessed whether conditioned medium from parental cells could restore proliferation in TS-KO cells when added to their growth medium. Surprisingly, however, this treatment was insufficient to re-establish proliferation of TS-KO cells (**Fig. 1K**). Similarly, parental cells seeded on a transwell-membrane, that was impermeable to mammalian cells but allowed sharing of medium with TS-KO cells, failed to restore growth of the TS-KOs (**Fig. 1L**), thereby validating a need for a physical connection between the cells sharing dTMP.

In summary, dTMP-deficient cells may attain proliferation by acquiring dTMP through direct contact with surrounding viable cells. These observations suggest the dispensability of *de novo* pyrimidine synthesis for growth. Importantly, we have identified a **collective mode of proliferation** relayed by intratumoral equilibration of dTMP.

### Collective proliferation drives pathophysiological features of cancer

Having established collective proliferation *in vitro* and in patients, global transcriptomic changes were analyzed by RNA sequencing in A549 to see how TS-KO cells transition from non-proliferative state to collective growth with parental cells. Differentially expressed genes (DEGs) were used to obtain gene signatures distinguishing non-proliferative TS-KO cells and parental controls (**Extended Fig. 6A**). To identify the molecular changes undertaken during transition from non-proliferative to proliferative state, TS-KO were collected from puromycin-treated coculture after growth restoration and compared to TS-KO cells cultured alone (**Extended Fig. 6B**). Finally, to understand the extent to which proliferative TS-KO cells regained the features of control cells, we compared proliferative TS-KO cells to parental controls (**Extended Fig. 6C**). Epithelial-mesenchymal transition (EMT), which is associated with metastasis and chemoresistance in cancer ^8^, was significantly enriched in non-proliferating TS-KO compared to TS-expressing controls (Extended Fig. 6A, **Extended Fig. 6D**). However, during transition to replicative state EMT signatures were downregulated in proliferative TS-KO cells compared to TS-KO cells with growth arrest (Extended Fig. 6B), while still maintaining higher EMT compared to parental controls (Extended Fig. 6C). Opposite trend was observed for the oncogenic KRAS signature, which was reduced in TS-KO cells, and was restored as TS-KO cells regained proliferation (Extended Fig. 6A-C). This implies that cells with dTMP deficiency differ significantly from cells that synthesize endogenous dTMP. Then, these signatures were investigated in *TYMS^-^TK1^-^* cells from the single cell transcriptomic data from patients (Extended Fig. 4). As in the proliferative TS-KO cells in coculture, KRAS, EMT signaling, and senescence-associated genes were upregulated in *TYMS^-^TK1^-^*cells whereas pyrimidine metabolism was downregulated (**Fig. 2A**). We then further explored the role of senescence, which contributes to many cancer phenotypes^9,10^. TS-KO cells were highly positive for the senescence makers β-galactosidase (**Fig. 2B, Extended Fig. 6E**) and p21 (**Extended Fig. 6F**). Based on the trends in the gene signatures from the TS-KO cells (Extended Fig. 6A-B), we speculated that they would at least partially revert senescence to regain proliferation in coculture. Therefore, we employed quantitative-image-based cytometry (QIBC) to quantify p21 protein expression in both TS-KO and control cells in cocultures at single-cell level. However, the mean expression of p21 in proliferative TS-KO cells in coculture remained unchanged compared to the TS-KO cells cultured alone (**Fig. 2C-D, Extended Fig. 6G**). Same observation was made when p21 levels were quantified by western blot (**Extended Fig. 6H**), indicating that cells in coculture could proliferate despite senescence markers, in line with previous observations^11^. RNA sequencing analysis also associated TS-KO cells with an upregulation of inflammatory signatures (Extended Fig. 6D). Since pyrimidine imbalance has been linked to cGAS-STING pathway, which is responsible for innate immunity response^12^, we quantified active cGAS in cytoplasmic micronuclei^13^ in dTMP deficient cells and found its activation to be strongly increased in both TS-KO and TS-TK1-KO cells compared to control cells (**Fig. 2E**). Further, protein quantification of phosphorylated STING and its downstream activator TBK-1 showed upregulation in dTMP deficient cell (**Fig. 2F, Extended Fig. 6I-J**). Since STING pathways is triggered by cytosolic DNA^14^, we quantified the levels of cytosolic single stranded DNA (ssDNA) in the cells after labelling them with CldU (5-Chloro-2’-deoxyuridine) and found high amount of ssDNA accumulated in TS-KO cells (**Fig. 2G**). Finally, the expression of γH2AX (a DNA-damage marker) was quantified, as nucleotide imbalance mediated cGAS-STING pathway has been previously associated with genome integrity^15^. Higher γH2AX levels were observed in both TS-KO and TS-TK1-KO cells compared to control cells (**Fig. 2H**). However, remarkably, donor parental cells in cocultures also showed significant increase in γH2AX activation in A549 (**Fig. 2I-J, Extended Fig. 6K**) and Calu-1 cells (**Extended Fig. 6L**), implying that dTMP insufficiency in the “donor” or the “acceptor” cells during collective proliferation could be sufficient to induce replicative stress and DNA damage.

**Figure 2.**
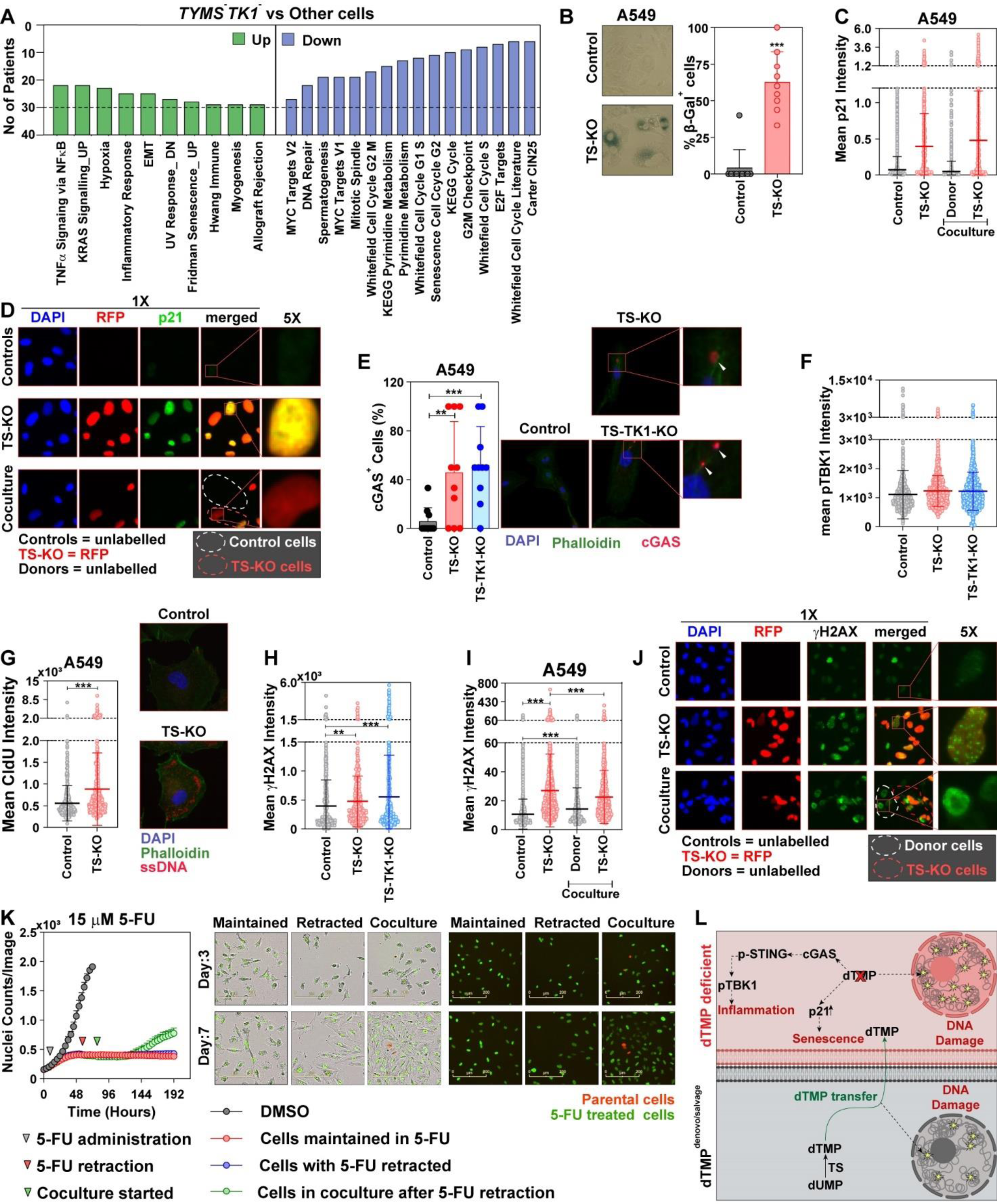
dTMP deficiency is associated with key pathophysiological features of cancer. **(A)** Pathways that are differentially regulated between *TYMS^-^TK1^-^* cells and other populations from cancer cells of NSCLC patients’ scRNA seq (Extended Fig 4). Only those pathways that were significantly regulated in more than 10 patients are included. **(B)** β-galactosidase staining in A549 TS-KO cells compared to controls, showing increased senescence in cells with TS-KO along with quantification of positive cells from the representative images. Bar represents mean ± SD of cells counted from 10 images; statistical analysis is Student’s t-Test. **(C-D**) Quantitative-image-based cytometry (QIBC) showing quantification of mean p21 expression (represented as mean intensity of immunofluorescent labelled p21 normalized to nucleus size) in A549 TS-KO cells with growth defect or in collective proliferation in coculture with parental A549 cells as donor (along with representative images). Each dot represents the mean intensity of p21 in a single cell, the longer line represents mean and shorter lines represent SD. Statistical analysis is Student’s t-Test. **(E)** Immunofluorescent staining of activated ssDNA-sensor cGAS in A549 cells with dTMP deficiency. Insets show cGAS signaling from cytoplasmic nuclear bodies, indicated with white arrows. Bars represent mean ± SD of cGAS^+^ cells counted from 10 images and normalized to the total cell counts/image. Statistical analysis is Student’s t-Test. **(F)** QIBC showing increase of p-TBK1, which is the effector protein of cGAS-STING pathway, significantly increased in cells with dTMP deficiency. Dots represent the mean intensity of p-TBK1/cells, the longer line represents mean and shorter lines represent SD. Statistical analysis is Student’s t-Test. Please see the representative images in the Extended Fig. 6I. **(G)** QIBC quantification of ssDNA in TS-KO cells quantified after staining with CldU. Dots represent the mean intensity of CldU/cell, the longer line represents mean and shorter lines represent SD. Statistical analysis is Student’s t-Test. **(H)** Quantitation of DNA damage marker γH2AX/cell in TS-KO A549 cells as quantified by QIBC. Dots represent cells, the longer line represents mean and shorter lines represent SD. Statistical analysis is Student’s t-Test. **(I-J)** Quantitation of γH2AX/cell in TS-KO A549 cells in coculture with donor parental cells, showing increased DNA damage in the donor parental cells in coculture compared to the controls. Dots represent cells, the longer line represents mean and shorter lines represent SD. Representative images and 5X zoom in the inset show increased γH2AX in parental cells marked by white broken lines. Statistical analysis is student’s t-Test. **(K)** Real-time proliferation of A549 cells treated with sublethal doses of 15 µM 5-FU (TS inhibitor), which ablates the growth of cells. Parental cells were added to the coculture after drug retraction showing that 5-FU induced growth arrest could be restored by coculture with parental cells. Data is mean ± SD of three replicates. Significance is calculated by two-Way ANOVA with Tukey’s multiple comparison. **(L)** Schematic representation of the pathophysiological features of dTMP deficient cells. Loss of dTMP leads to activation of DNA damage, senescence via p21 activation, and inflammation via cGAS-STING pathway. During collective proliferation, dTMP is passed from donor parental cells to dTMP deficient cells, creating DNA damage in the donor cells.

Finally, since dTMP depletion is used as an intervention strategy in clinical oncology, we investigated if collective proliferation might render resistance to drugs that target TS. Therefore, we treated A549 cells with sublethal doses of the TS inhibitor 5-flurouracil (5-FU), which abrogated growth without killing. After drug retraction, parental cells were added to 5-FU treated cancer cells and were found to significantly accelerate their growth restoration, indicating that collective proliferation could also contribute to chemoresistance (**Fig. 2K, Extended Fig. 6M**). To conclude, collective proliferation drives several physio-pathological features of cancer by facilitating the proliferation of dTMP-deficient (including chemotherapy-treated) cells without altering their p21 status and by creating a nucleotide deficit that propagates genomic instability within the cancer population (**Fig. 2L**).

### Tumor microenvironment (TME) cells contribute to collective proliferation

Next, we wanted to evaluate the stringency of the pairing of donor and dTMP-deficient cells in collective proliferation. Therefore, we attempted growth rescue of TS-KO A549 cells (of adenocarcinoma origin) with different lung epithelial cell lines (Calu1 of squamous cell carcinoma origin and BEAS-2B of non-cancerous origin). Remarkably, both Calu-1 and BEAS2B rescued the growth of A549 TS-KO cells in coculture (**Figure 3A**). This observation prompted us to extend our investigation to non-epithelial cells previously associated with nucleoside secretion^16-18^. Of these, tissue-resident fibroblasts (**Figure 3B**), cancer-associated fibroblasts (CAFs, **Figure 3C**) and peripheral blood mononuclear cells (PBMCs) (**Figure 3D**), as well as the transformed Jurkat cells of T-cell origin (**Figure 3E**), failed to restore growth of TS-KO cells. However, human umbilical vein endothelial cells (HUVEC) cells and THP-1 monocytes restored proliferation of TS-KO cells (**Figure 3F-G**). Further looking into macrophages polarization states, (**Figure 3H**), M0 and M2 macrophages rescued the growth defect in TS-KO cells, whereas M1 macrophages did not (**Figure 3I**). Considering that protumor M2 macrophages support TS-KO cell growth and that cancer cells polarize macrophages to a M2-like phenotype (tumor-educated macrophages, TEMs) to secrete pyrimidines^19^, we administered A549 cell culture medium (CM) to M0 macrophages to polarize them to TEMs. CM was then collected from the TEMs and supplemented with fresh cell-culture medium for TS-KO cells. While TEMs had a positive effect on TS-KO proliferation when co-cultured (like M2 macrophages), CM from the same TEMs did not facilitate TS-KO cell proliferation (**Extended Fig. 7A-B**), further confirming that dTMP sharing involves non-secretory routes. We then analyzed scRNA-sequencing data from patients and found a positive correlation between endothelial cells and number of *TYMS^-^TK1^-^* cells, but not with M2 macrophages and epithelial cells (**Figure 3J-L, Extended Fig. 7C**). We further performed a neighborhood analysis and found an increased propensity for endothelial cells in *TYMS^-^TK1^-^* cells (**Figure 3M, Extended Fig. 7D**). This could imply that different TME donors could be employed in different tissues depending on the proximity and availability. Thereby a non-specific nature of collective proliferation emerges, where dTMP deficient cancer cells can make physical contacts with different types of TME cells to maintain growth.

**Figure 3.**
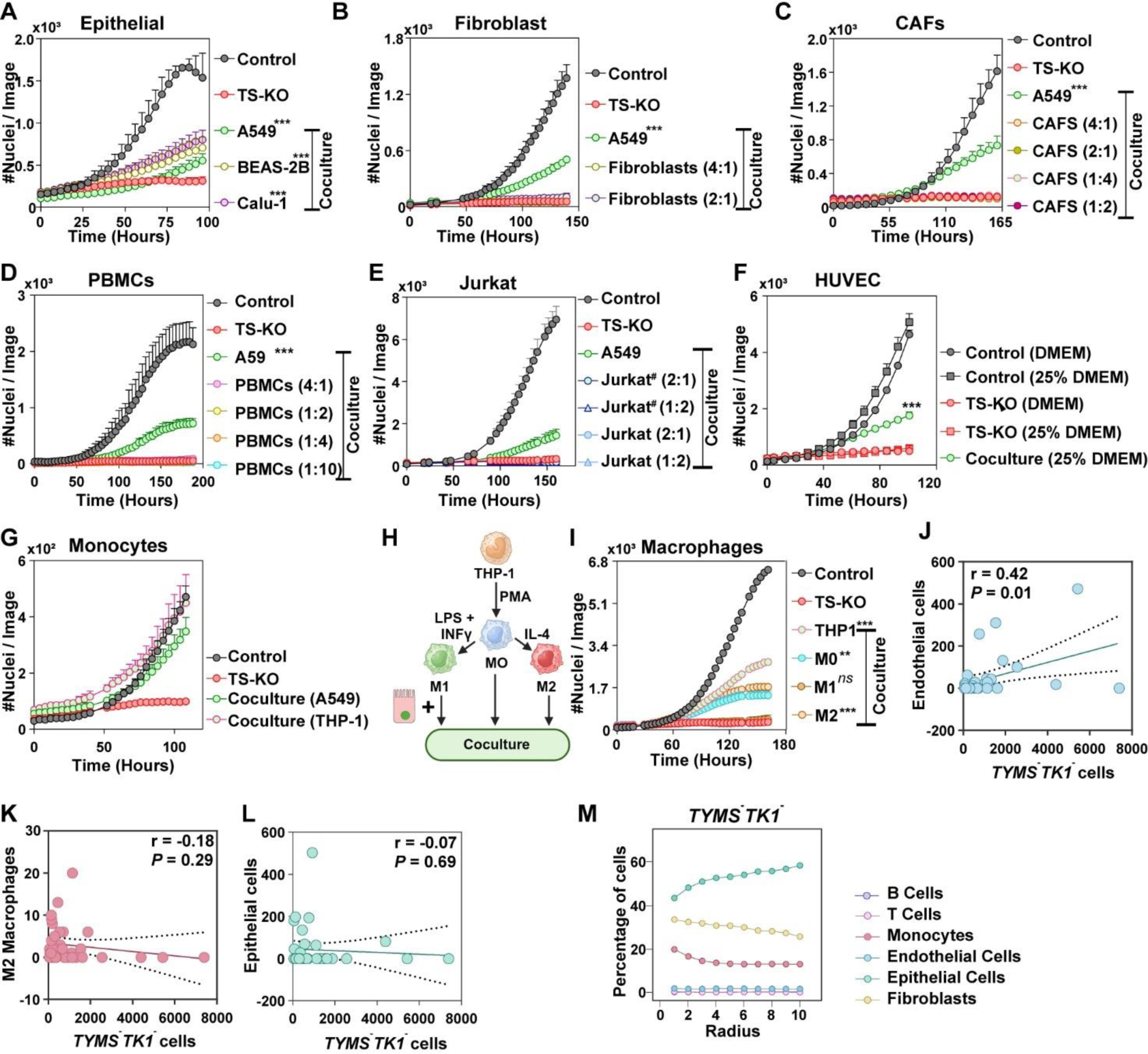
The contribution of tumor microenvironmental (TME) cells to collective proliferation. **(A)** Proliferation kinetics of A549 TS-KO in coculture with Calu1 and BEAS2B, showing that collective proliferation can be coordinated between a LUSC cell line (Calu-1), a non-cancerous cell line of epithelial origin (BEAS2B), and an LUAD cell line (A549). **(B-C)** Coculture of A549 TS-KO cells with normal fibroblast cells and cancer associated fibroblasts (CAFs). **(D)** A549 TS-KO cells cocultured with peripheral blood mononuclear cell (PBMCs), that is highly enriched in lymphocytes. **(E)** Proliferation curve showing inability of either activated (Jurkat^#^) or activated Jurkat cells to re-establish growth of A549 TS-KO cells. **(F)** Proliferation curve showing human umbilical vein endothelial cells (HUVEC) supporting collective proliferation of A549 TS-KO cells. **(G)** A549 TS-KO cells undergoing collective proliferation in presence of THP-1 cells of monocytic lineage. **(H)** A scheme showing induction of THP-1 monocytes to macrophage with different polarization. **(I)** Coculture of A549 TS-KO with unpolarized M0 macrophages, M1 macrophages, or M2 macrophages lineages from (H), showing collective proliferation being supported by all macrophages except the M1 type. **(J-L)** Correlation between number of different TME cells (endothelial cells, M2 macrophages and epithelial cells) and *TYMS^-^TK1^-^* cells from scRNA data from Extended Fig. 4. Correlation is calculated by Pearson’s rank correlation. **(M)** Neighborhood analysis showing increased propensity of *TYMS^-^TK1^-^* cells to be in neighborhood of epithelial cells. In experiments (A-I) data points represent mean ± SD for three replicates and the statistical test is Two-Way ANOVA with Tukey’s multiple test.

### dTMP exchange is mediated through gap junctions

Having established that physical contact between cells is necessary for collective proliferation, the next step was to investigate the molecular nature of this coupling. RNA sequencing showed that *GJA1* and *GJD3* were upregulated in growth-deficient and proliferative TS-KO cells, compared to parental control cells (**Extended Fig. 8A**). *GJA1* and *GJD3* code for proteins Connexin43 (Cx43) and Connexin30.2 (Cx30.2), respectively. Cx43 and Cx30.2 can hexamerize to form gap junctions between cells, which generates a syncytium for intercellular metabolite exchange and may underpin the growth rescue effect^20,21^. Theorizing that TS-KO cells supply dTMP via gap junctions in coculture settings, the possibility that these cells are diffusively coupled was tested by fluorescence recovery after photobleaching (FRAP). Specifically, a monolayer of NSCLC cells was stained with calcein red-orange AM (size comparable to thymidylate), and a single cell randomly selected in a cluster was bleached within its limits. The post-bleaching recovery of fluorescence was recorded in the bleached cell. Results indicated a strong recovery that was blocked in the presence of carbenoxolone (CBX), a pan-gap junction inhibitor (**Fig. 4A, Extended Fig. 8B**). This implied that gap junctions are potential conduit for the diffusion of dTMP between cells. This was demonstrated by treatment of cocultures with CBX, which significantly reduced the growth rescue of the TS-KO cells at a concentration of CBX that had no effect on the proliferation of the parental cells (**Fig. 4B, Extended Fig. 8C-D**). In contrast, inhibition of nanotunnels (that form another mode of intercellular communication) by SMIFH2^22^ did not reduce TS-KO cell growth in coculture (**Extended Fig. 8E-F**). To further validate the role of gap-junctions in three-dimensional setting, we generated spheroids containing TS-KO and parental control cells treated with CBX or vehicle control. While TS-KO cells proliferated within spheroids containing parental cells, their growth was inhibited when these cocultures were treated with CBX (**Fig. 4C, Extended Fig. 8G**). Similarly, CBX treatment prevented THP-1 monocytes from restoring TS-KO cell growth (**Fig. 4D**). Together, these data confirm gap junction-mediated cell coupling as the route for dTMP transfer.

**Figure 4.**
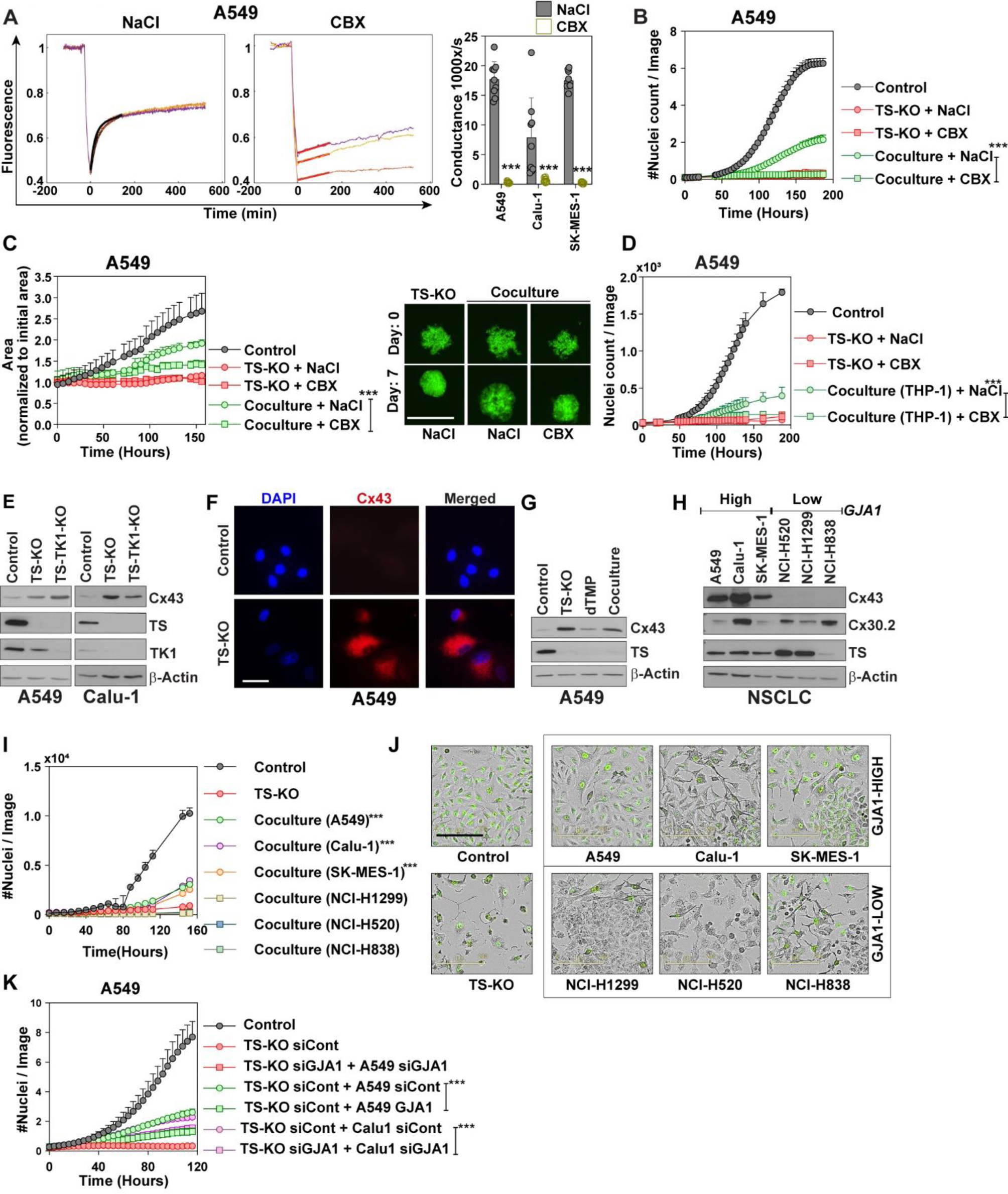
Gap-junctions (GJs) mediate dTMP sharing. **(A)** Fluorescence recovery after photobleaching (FRAP) showing connectivity of indicated cells, measured by their ability to reconstitute fluorescent dye after bleaching, in presence of either NaCl or 100 μM of the gap junction inhibitor Carbenoxolone (CBX). Each bar represents the mean ± SD of 10 replicates. Significance is determined using One-Way ANOVA analysis. Fluorescence recovery for for Calu-1 and SK-MES-1 cells are shown in Extended Fig. 8B. **(B)** Incucyte proliferation assay measuring growth of A549 TS-KO cells in coculture with parental A549 cells in presence of either 100 μM CBX or NaCl. Significance is determined using a Two-Way ANOVA analysis with Tukey’s multiple comparison. **(C)** Collective growth of TS-KO cells, as grown in 3D-spheroids, in presence of either 100 μM CBX or NaCl. Images for spheroids showing GFP signal, scale bar represents 800 μm. Each point represents the mean ± SD of five replicates. Significance is determined using a Two-Way ANOVA analysis with Tukey’s multiple comparison. See representative images of control and TS-KO spheroids in Extended Fig. 8G. **(D)** Collective growth of TS-KO cells in presence of THP-1 monocytes abrogated by supplementation of 100 μM CBX compared to NaCl. The data are mean ± SD of 3 replicates. Significance is determined using a Two-Way ANOVA analysis with Tukey’s multiple comparison. **(E)** Increased expression of Cx43 proteins, which hexamerizes to form GJA1 quantified in TS-KO and TS-TK1 KO cells. **(F)** Immunofluorescence images showing increase of Cx43 (the protein coded by GJA1) in TS-KO cells. Scale bar represents 200 μm **(G)** Quantification of Cx43 in proliferative A549 TS-KO cells compared to the controls and non-proliferative TS-KO cells. **(H)** NSCLC cell lines classified high and low based on the expression of *GJA1* (Cx43). **(I)** The inability of *GJA1* low cells to promote collective growth in TS-KO cells, as quantified by Incucyte S3. The data are mean ± SD of three replicates. Significance is determined using a Two-Way ANOVA analysis with Tukey’s multiple comparison. Significance is compared to the TS-KO cells. **(J)** Representative images from (I) showing that TS-KO cells failed to restore their normal morphology when they were cocultured with *GJA1*-Low cells. Scale bar represents 200 μm. **(K)** Simultaneous siRNA (80 nM) mediated knockdown of *GJA1* in A549 TS-KO and parental cells, and its abrogative effect on collective growth. The data are mean ± SD of three replicates. Significance is Two-Way ANOVA with Tukey’s multiple comparison.

Since RNA sequencing data identified *GJA1* and *GJD3* as the main candidates for forming connexins, their protein levels were quantified in TS-KO cells. Expression of Cx43 (encoded by GJA1) was significantly increased in dTMP-deficient TS-KO and TS-TK1-KO cells, compared to the parental cells, as confirmed by western blots (**Figure 4E**) and Cx43 immunofluorescence staining (**Figure 4F, Extended Fig. 9A**). Further, Cx43 expression was quantified in proliferating TS-KO cells by western blot and was found to be comparable to that in non-proliferating TS-KO cells (**Figure 4G**). Immunofluorescent staining of donor A549 cells from coculture with TS-KO showed that the parental cells also increased Cx43 expression (**Extended Fig. 9B**). Therefore, Cx43 was postulated to be the key connexin involved in dTMP equilibration. This was further confirmed in computation based autodocking of dTMP to GJA1 protein, which predicted binding of dTMP to open conformation of Cx43 (**Extended Fig. 9C-D**). Consequently, the absence of Cx43-mediated coupling was hypothesized to abolish the proliferation of TS-KO cells in coculture. Therefore, based on *GJA1* mRNA expression (**Figure 9E**), cells with undetectable levels of *GJA1* were identified for further studies. After confirming the lack of Cx43 expression by western blot (**Figure 4H**), NCI-H1299, NCI-H520 and NCI-H838 cells were cocultured with TS-KO cells. While A549, Calu-1, and SK-MES-1 (Cx43 positive cells) restored growth of the TS-KO cells, the cells lacking detectable Cx43 (NCI-H1299, NCI-H530 and NCI-H838) failed (**Figure 4I**). Moreover, while coculture with Cx43 expressing cell lines reverted TS-KO cells to normal morphology, coculture with GJA1-low cells showed TS-KO cells with stressed morphology (**Fig. 4J**). To further validate the role of GJA1, siRNA-mediated knockdown of GJA1 was introduced into TS-KO cells, as well as in A549 and Calu-1 cells that were used as dTMP donors. While GJA1 knockdown did not affect parental cell growth (**Figure S9F-G**), it significantly reduced the growth of TS-KO cells cocultured with A549 or Calu-1 cells (**Figure 3K**). Altogether, these findings confirm GJA1 as the major channel of dTMP equilibration.

### Gap junction-dependent *Tyms*^-^*Tk1*^-^ tumor growth *in vivo*

To test whether cells lacking *de novo* and salvage dTMP synthesis can form tumors *in vivo*, we knocked out *TYMS* and *TK1* in A549 cells (**Extended Fig. 10A-B**) and injected them in the flanks of NSG mice. Despite being unable to grow *in vitro* in the absence of parental dTMP donors (**Extended Fig. 10C**), TYMS-TK1-KO cells formed tumors in mice (**Figure 5A**). Protein quantification of isolated tumors confirmed the lack of dTMP synthesizing enzymes together with a significant increase in Cx43 (expressed by GJA1) in TS-TK1-KO tumors (**Figure 5B)**. This result prompted generation of a lung cancer mouse model with genetic ablation of *Tyms* and *Tk1*. Therefore, lung-tropic adeno-associated viruses (AAV) carrying gRNAs for humanized *Kras^G12D^* mutation, *p53*, *Tyms*, and *Tk1* were delivered to C57BL/6 ROSA-LSL-Cas9 mice^23^ to obtain the *Kras^G12D^p53*^-/-^*Tyms*^-/-^*Tk1*^-/-^ (KPTT) model (**Figure 5C, Extended Fig. 10D)**. Mice with *Kras^G12D^p53*^-/-^ (KP) derived tumors^24^ were used as controls. Similar to the results from NSG mice, we found that tumors could form and grow in mice without Ts and Tk1 (**Fig. 5D-E**). However, the cells derived from a KPTT tumor failed to grow *in vitro* without dTMP supplementation and displayed a senescent morphological phenotype upon dTMP deprivation (**Fig. 5F**), similar to observations in human cells. These findings indicate that *in vivo* their growth depends on an exogenous source of dTMP. Furthermore KPTT-derived cells (that could not grow without dTMP) could form new tumors when injected in the flanks of wild type C57BL/6 mice (**Fig. 5G**). This confirms their tumorigenicity despite the lack of intrinsic dTMP synthesis.

**Figure 5.**
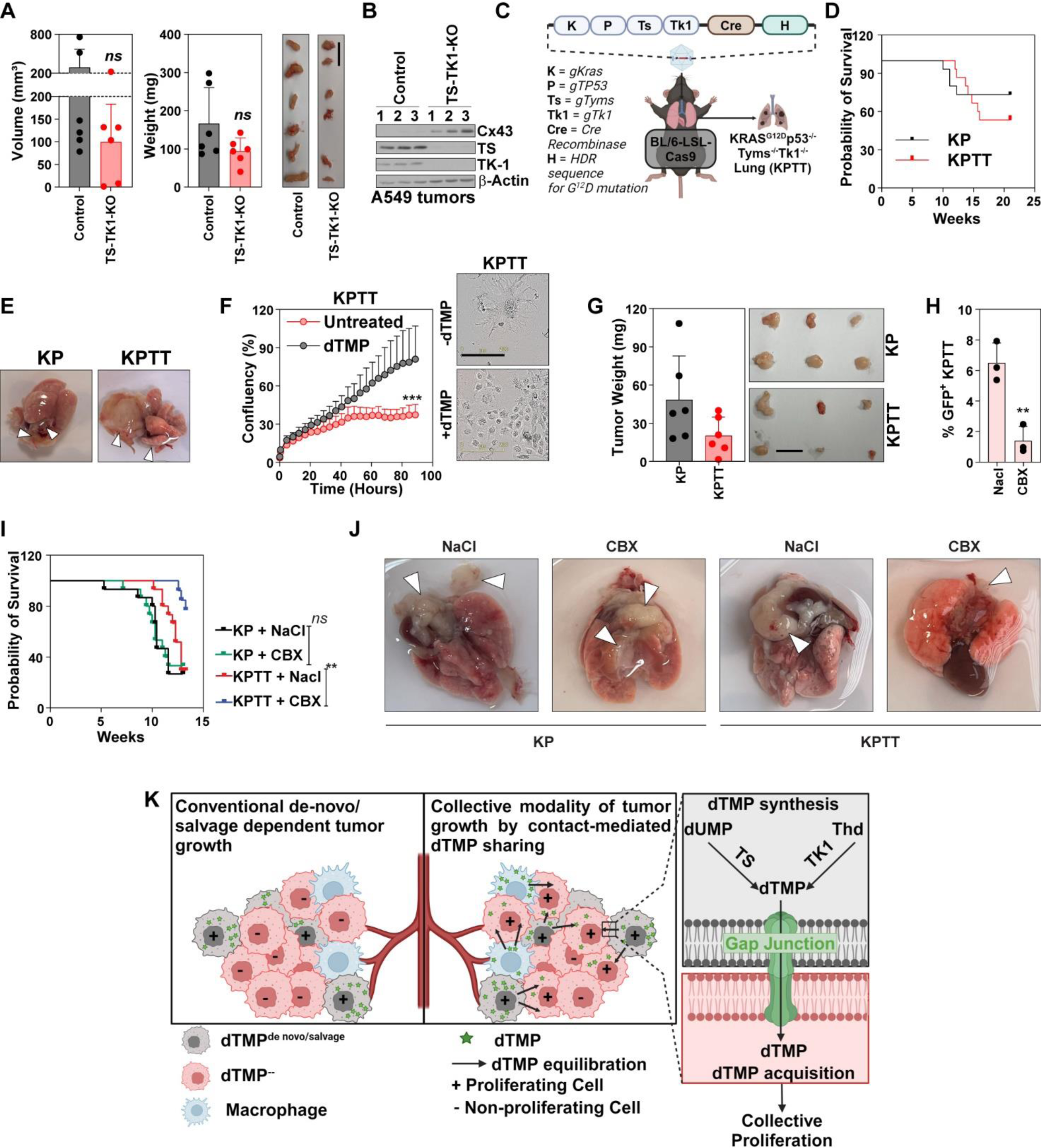
Gap junction dependent growth of dTMP-deficient growth of NSCLC tumors. **(A)** Weights and volumes of tumor formed from A549 TS-TK1-KO cells in flanks of NSG mice along with the pictures of excised tumors. TS-TK1-KO tumors were excised five days after the controls. Bars are mean ± SD of tumors from 6 flanks of 3 mice and significance is calculated by Student’s t-Test. Scale bar represents 1 cm. **(B)** Westen blot quantification of TS and TK1 in tumors excised from (A) showing depletion of TS and TK1 proteins and increased expression of Cx43 (*GJA1*) in TS-TK1-KO cells**. (C)** C57BL/6 expressing Cas9 were intubated with AAV9 vectors carrying gRNA for *Kras* (along with G12D HDR sequence), *p53*, *Tyms,* and *Tk1* resulting in lung specific mutations in targeted genes. Intubated mice develop Kras^G12D^ and p53^-/-^ driven tumors that lack expression of *Tyms* and *Tk1* genes, resulting in KPTT tumors. **(D)** Comparison of tumors formed by mice harboring tumor specific mutation in lungs of C57BL/6 mice. KP (Kras^G12D^ p53^-/-^) mice have been used as controls. **(E)** Representative images from lungs with tumors from the mice in (C). **(F)** Cell line established from KPTT tumor dependent on exogenous dTMP supplementation (10 μM) for proliferation and undergoing senescense-like enlarged phenotype in absence of dTMP. The data are mean ± SD of three replicates. Significance is Two-Way ANOVA with Tukey’s multiple comparison. **(G)** Weight of tumors excised from subcutaneous implantation of KP and KPTT tumors in C57BL/6 along with representative images. Each bar represents the mean ± SD of tumors from 6 flanks of 3 mice. Significance is calculated by Student’s t-Test. Scale bar represents 1 cm **(H)** Flow cytometry quantification of GFP^+^ KPTT cells cocultured with syngeneic Ladi 2.1 cells derived from a lung tumor of a C57BL/6 mouse. Cocultures were treated either with 100 μM CBX or the vehicle control. The bars are mean ± SD of three replicates and significance is calculated by Student’s t-Test. **(I)** Survival of KPTT mice compared to KP mice when treated with 40 mg/kg CBX or corresponding volume of the vehicle control NaCl. (n = 15 for all the groups except the KPTT + CBX where n = 14). Statistic is calculated by Log-Rank (Mantel Cox) test. **(J)** Images of lung with tumors from (H). KPTT+CBX lungs are from the same week of the representative KPTT + NaCl lung, at the time of the termination of experiment. **(K)** The current model of proliferation relies on activation of canonical *de novo* and/or salvage pathways in each proliferating cell, whereas the model of collective proliferation postulates that growth is a cumulative sum of proliferation of cells with *de novo* and salvage and cells that rely of dTMP sharing via gap junctions.

Next, to test the involvement of gap junction in the growth of KPTT cells, the cells were cocultured with a syngeneic cell line Ladi 2.1 (derived from a lung tumor of a C57BL/6 mice) in the presence and absence of CBX. Since KPTT tumors express cytoplasmic GFP^23^, the fluorescence signal was used as the proliferation readout. KPTT cells grew in coculture with Ladi 2.1 cells but proliferation was significantly reduced in the presence of CBX (**Figure 5H, Extended Fig. 10E**), indicating the role the of gap junctions in their growth. To test this further *in vivo*, mice were intubated to generate KP and KPTT tumors, followed by subcutaneous injection of CBX or vehicle control (NaCl) for 8 weeks. Although no difference was observed in the survival of CBX treated KP mice, the drug significantly reduced and delayed the formation of tumors in KPTT mice and increased their mean survival (**Fig. 5I**). Compared to vehicle control, in the CBX treated KPTT group three mice formed tumors comparable to controls and three mice formed small tumors which didn’t develop symptoms that would meet the humane endpoint criteria (**Fig. 5J**). The other mice from this group didn’t form detectable tumors. H&E staining of mouse lungs showed that CBX-treated KPTT mice developed tumors of comparatively smaller size with visibly intact parenchyma, compared to the larger tumor burden in controls (**Extended Fig. 10F**). Taken together, these results confirm that tumor growth *in vivo* can proceed without active dTMP synthesis, facilitated by gap junction-mediated dTMP acquisition from surrounding tissue.

Our data presents a strong rational to revisit the canonical notion that synthesis of dTMP in every single transformed cell drives tumor growth and establish a new paradigm of collective proliferation. In the postulated model proliferation can be maintained by dTMP being synthesized in some cells within a tumor (both of cancer and non-cancer origin) and redistributed to other cancer cells via gap junctions (**Fig. 5K**).

## Discussion

The crosstalk of tumor entities at multiple levels is fundamental to cancer growth, such that immunotherapeutic abrogation of receptor-driven interaction between cancer cells and immune cells has been one of the major breakthroughs in the clinical intervention of NSCLC and other cancers^25,26^. Metabolic codependency and cross-feeding of metabolites between TME and cancer cells further adds to this complexity and is equally important for tumor growth^27,28^. By breaking the dogma of the indispensability of cancer-specific dTMP biosynthesis and by identifying dTMP equilibration within tumor as an important growth mechanism, this study advances the concept and postulates a model of collective proliferation.

The concept of collective proliferation repurposes metabolite sharing and resource redistribution as an energy-efficient compensatory mechanism for metabolic vulnerabilities in tumors^29^. In particular, the redistribution of thymidylate within tumors might be a functional rheostat for adapting to metabolic heterogeneity^30^, overcoming the challenges of the harsh growth environment^31^ and modulating the dynamics of TME^32^. Moreover, since proliferation is a fundamental physiological process, the concept of collective proliferation may have relevance not only in other cancers, but also in embryogenesis, tissues regeneration, and other proliferative diseases (atherosclerosis, idiopathic pulmonary fibrosis, scleroderma, etc.). Therefore, it is pertinent to understand all the aspects of collective proliferation in more extensive and dedicated studies, which includes the central role of dTMP in collective proliferation, origin and characterization of cells that lack dTMP deficiency, interaction of dTMP deficient cells with donor cells, effect on therapy response, and finally how dTMP sharing contributes to the evolving conundrum of pyrimidine metabolism in cancer.

Firstly, our data that dTMP is the only nucleotide shared during collective proliferation are very remarkable. dTMP is distinct as it is the nucleotide moiety exclusive to the dNTP pool, suggesting it might possess characteristics that are not shared by other dNTPs. These unique features might have important roles in genomic instability and DNA repair during xenobiotic stress, as it has previously been shown that thymidine metabolism is an important determinant of telomere length^33^.

Further, understanding the origin of dTMP deficient cells during clonal expansion will be a vital addition to proliferation dynamics in tumor growth. The prime question is whether these cells are reminiscent of cells that fail to overcome oncogene-induced senescense^34^ or they are more plastic to modulate dTMP synthesis to the external cues.

This study links the lack of dTMP expressing genes with certain key pathological features. However, a detailed characterization of dTMP deficient cells will unravel the biological nature and clinical implications of this association. Of note, it is important to understand the fate of senescent cancer cells that rely on exogenous dTMP supply, as senescence plays a critical role in cancer progression and treatment response^35^. Moreover, the observed connection with cGAS-STING pathway, a cellular detector of cytoplasmic DNA and innate immune response activator, suggests that dTMP availability (and its fluctuations) may influence tumor inflammation.

Furthermore, it is important to note that collective proliferation is an interactive process and includes cells from different lineages, requiring a thorough exploration of its dynamic nature. A detailed study is needed to understand the association between dTMP deficiency and activation of connexin proteins, especially in the donor cells, as we observed that dTMP-deficient cells increased the expression of GJA1 in cells that already had detectable levels of GJA1, while failing to induce its expression in other cells where GJA1 was initially undetectable. Therefore, more detailed insight into epigenetic modulation of these interactions is required. In addition, one of the most intriguing observations that calls for further exploration is the induction of DNA damage in donor cells that participate in dTMP equilibration. This could indicate that collective proliferation is a mean by which genomic instability is propagated in tumors and could have different impacts the non-cancerous donor cells (such as monocytes and endothelial cells) depending on the context of their interaction with cancer cells.

A major implication of the outcomes of this research relates to therapeutics for NSCLC and other cancers, since targeted metabolic perturbation is central to chemotherapy, including TS targeting drugs like 5-flurouracil and pemetrexed^36^. In this context, metabolic redistribution by refractory cells might represent a so far overlooked mechanisms of chemoresistance to anti-dTMP-based drugs such as 5-flurouracil and pemetrexed^37,38^. Collective proliferation could also contribute to resistance against immune therapy where dTMP deficient cells act as nucleotide sponges that affect the differentiation and exhaustion of immune cells^39^. Subsequently, the model may refine therapeutic avenues, for instance by repurposing CBX, which is approved for peptic, esophageal, and oral ulceration^40^, or other gap junction inhibitors, for clinical testing of combination therapies.

Finally, collective proliferation, particularly in the context of dTMP equilibration, adds another dimension to the evolving complexity of pyrimidine metabolism in cancer. Recent advances have attributed moonlighting functions to classical metabolites^41,42^. This also includes pyrimidine metabolites that can play a critical role in differentiation^43-47^, act as a cofactor in lipogenesis^48^, and intercellularly feed glycolysis under metabolic stress^49-51^ beyond their role in DNA synthesis and proliferation. The concept of dTMP sharing in context of collective proliferation highlights the importance of pyrimidine metabolism beyond the terminal outcome of metabolic alterations in cancer to support cell proliferation. Therefore, its critical and diverse roles in survival and growth necessitate studies to identify the cell intrinsic and extrinsic cues that direct pyrimidine metabolites to feed into other essential pathways, compensate for intercellular nutritional stress, or support proliferation of adjacent cells that lack dTMP synthesis.

In conclusion, this study repositions proliferation in the context of metabolic resource sharing and establishes a more dynamic and plastic nature of tumor growth. Further exploration of this mechanism will not only enhance the current understanding of the interplay among tumor growth, heterogeneity, and pathology but will also provide a new perspective to target proliferation.

## Acknowledgements.

The authors wish to acknowledge the support of Metabolomics Core Facility, EMBL, Heidelberg, Germany for their support with mass-spectrometric quantification, and Dr Thomas Solomon at Blazon Scientific for assistance with drafting the manuscript. The study was funded by Hallas-Møller Ascending Investigator Grant from Novo Nordisk Foundation (PC), Independent Research Fund Denmark 0134-00012B (PC), Lundbeck Foundation, Denmark, R380-2021-1264 (MAS), International Association for the Study of Lung Cancer (MAS), Danish Cancer Society (AMAR), Danish Cancer Society (AMT). Illustrations were drawn using Biorender.

## Author Contribution

P.C., M.A.S. and A.M.A.R were involved in project conceptualization and development and the preparation of the manuscript. P.C. was involved in supervision. M.A.S., A.M.A.R., L.P., S.V., S.S.M., P.N.G. A.M.T. and L.B.G conducted the experiments. V.R. and N.P.P. conducted the bioinformatic analysis. L.C.B and M.Ve. performed molecular modeling simulation. S.K., F.N., M.Vo and P.S. were involved in data acquisition.

## Materials availability

All the plasmids generated in this study are available from the lead contact without restriction or with a completed materials transfer agreement, whichever applies.

## Data and code availability

RNA sequencing data has been deposited to GEO dataset (GSE271721). Any additional information required to reanalyze the data reported in this paper is available from the lead contact upon request.

## Methods

### Cell lines

Lung cancer cell lines (A549, Calu-1, SK-MES-1, NCI-H1299, NCI-520, NC-H838) and THP-1 monocytes were purchased from ATCC. Normal fibroblasts and Cancer Associated Fibroblasts were provided by Dr. Pamela L. Strissel, at the University Clinic, Erlangen, Germany. Ladi 2.1 cells were obtained from Dr. Markus E. Diefenbacher, Biochemistry and Molecular Biology Biocenter, University of Würzburg, Germany. HUVEC cells were provided by Professor Ditte Caroline Andersen, Research Unit for Clinical Biochemistry, Odense University Hospital, Denmark. HEK-293T cells were obtained from Dr. Heiko Wurdak, Faculty of Medicine and Health, School of Medicine, University of Leeds, UK. The KPTT cell line was established from C57BL/6 mouse with genetic knock of Tyms and Tk1. All cells were cultured in DMEM supplemented with 2 mM of L-glutamine (Gibco), 100 U/ml of Penicillin–Streptomycin (Gibco), and 10% Fetal bovine serum (Gibco), except for THP-1 and Jurkat cells that were cultured in RPMI 1640 (Gibco) supplemented with 10% FBS (Merch) and 1% penicillin-streptomycin (Gibco). HUVEC cells were cultured in Endothelial Cell Growth Medium 2 medium (PromoCell) supplemented with the ready-made Supplement Mix. All cell lines were authenticated by STR profiling and tested regularly for mycoplasma contamination, kept in culture for no more than 20 passages and maintained at 37°C with 5% CO_2_ level in a humidified incubator.

### Lentiviral transduction

To produce lentiviral particles, HEK 293T cells were transfected with 8 µg expression vectors and 2 µg of third-generation lentiviral packaging system in complex with 24µg PEI (Polysciences) in 1.2 ml of 0.9% NaCl. After 48 hours, the supernatant containing the lentiviral particles carrying the gRNAs was collected, centrifuged, and filtered. For transduction, 50,000 cells were seeded in a 6-well plate and transduced with lentiviral vectors carrying transgenes in presence 32 μg polybrene (Sigma). Viruses were removed 24 hours post the transduction, and selection was started 48 hours post transductions. TS-KO cells and TS-TK1-KO cells were selected with 3 μg/ml puromycin (Sigma). Cells transfected with gRNA for TK1, DTYMK, and RRM1 were selected with 1 mg /ml hygromycin (Sigma). H2B-GFP or H2B-RFP transduced cells were flow-sorted for green and red positivity respectively. Plasmids and gRNA sequences are listed as Extended Table.

### Incucyte real-time Proliferation Assay

For Incucyte real-time proliferation assay, cells were seeded in 96-well plates at a density of 1,800 – 2,000cells/well. For cocultures 1,800 - 2,000 TS-KO or TS-TK-KO cells were seeded with donor cells in ratios mentioned in the respective figure legends. Plates were scanned at regular intervals of 4 hours – 24 hours using basic analyzer module. Incucyte 2021C software was used for analysis. In brief, 8 – 10 images were used to train the software to differentiate cells from background to calculate confluency as the area of the plate occupied by cells. Similarly, Top Hat background subtraction method was applied to the fluorescent signals to define nuclei. Processing definition was created from the training set that was applied to all scans and proliferation kinetics were created as proliferation over time and number of nuclei over time.

### Metabolite treatments

For metabolite treatments, H2B-GFP^+^ TS-KO cells were seeded at a density of 1800–2000 cells/well in a 96-well plate. Medium was changed on the following day and cells were then treated with metabolites as indicated in the figure legends. Plates were scanned and analyzed using Incucyte real-time proliferation assay. Thymidylate (dTMP), thymidine (Thd), deoxyuridine monophosphate (dUMP), and deoxythymidine triphosphate (dTMP) were purchased at Sigma. Deoxyadenosine triphosphate (dATP), deoxyguanosine triphosphate (dGTP), and deoxycytidine triphosphate (dCTP) were purchased at Promega.

### THP-1 differentiation

THP-1 monocytes were differentiated into macrophages by stimulation with phorbol 12-myristate 13-acetate (PMA). Briefly, 800,000 THP-1 cells were stimulated with 100 ng/ml PMA for two days in a 6-well plate. During the stimulation, the THP-1 cells would adhere to the surface of the well. To further differentiate the macrophages into M1- or M2 macrophages, the cells were stimulated with 100 ng/ml lipopolysaccharide (LPS) (Sigma-Aldrich) and 20 ng/ml interferon gamma (IFNγ) (Peprotech) or 20 ng/ml interleukin 4 (IL4) (Peprotech) respectively for 72 hours. During the M1- and M2 polarization, M0 macrophages were cultured in RPMI supplemented with 10% FBS and 1% penicillin-streptomycin.

### PBMC isolation

Peripheral blood mononuclear cells (PBMCs) were isolated from healthy individuals using Lymphoprep separation (StemCell Technologies) according to the manufacturer’s instructions. Isolated PBMCs were frozen in FBS with 10% DMSO. Prior to use in co-culture experiments, PBMCs were thawed and seeded in X-vivo medium (Lonza) supplemented with 5% human serum (Sigma) at a density of 1.5 x 10^6^ cells/well in a 24-well plate. The cells were rested for overnight in a 37°C, 5% CO2 incubator. To ensure an enrichment of T cells, cells were then collected from the 24-well plate by resuspension, eliminating cells that had adhered to the bottom of the well overnight.

### Coculture experiments

For coculture experiments, 1,800–2,000 H2B-GFP-TS-KO (or TS-TK1-KO cells) cells were seeded with dTMP donor cells in 96-well plates. Coculture was optimized by trying different ratios between TS-KO and the dTMP donor cells. For coculture with A549, Calu-1, SK-MES-1, BEAS-2B, NCI-H1299, NCI-H838, and NCI-520 a ratio of 1:4 was used. Jurkat cells were stimulated with CD3/CD28 dynabeads (Gibco) according to the manufacture’s protocol (Gibco) during the duration of the coculture. HUVEC cells were cocultured with TS-KO cells in presence of combined medium (25% DMEM: 75% HUVEC medium). The proliferation of the H2B-expressing cells was monitored using Incucyte S3.

### Tumor-educated macrophages

Tumor-educated macrophages (TEM) were generated as described previously^19^. Initially, 1.5 million A549 cells were cultured in DMEM supplemented with 10% FBS, 1% P/S, and 1% L-glutamine in a 10-cm dish. One day after seeding, the medium was changed on the 50% confluent A549 cells, and the cells were cultured for an additional 48 hours. Next, the conditioned medium was collected from the A549 cells and centrifuged at 300 × g for 5 minutes. The conditioned medium was then filtered through a 0.45 um filter and heat-inactivated for 15 minutes at 100°C. DMEM supplemented with 10% FBS, 1% P/S, and 1% L-glutamine was added to the conditioned medium to restore the initial volume. Like the *THP-1 differentiation* described above, 1.6 million THP-1 cells per well of a 6-well plate were stimulated with 100 ng/ml PMA for 48 hours. Then the medium was removed and replaced with 75% heat-inactivated conditioned medium from the A549 cells. The THP-1 cells were cultured for an additional 48 hours to polarize them into TEM. Then the conditioned medium from the TEMs was collected, centrifuged at 300 × g for 5 minutes, and heat-inactivated at 100°C for 15 minutes. Condition medium was added to was added to 1,800 A549 GFP^+^ TS-KO cells in three different ratios (1:4, 1:1 and 4:1) in a 96-well plate. The TEM macrophages were trypsinized and cocultured with KO cells. Proliferation was monitored using Incucyte S3 real-time proliferation assay.

### Transwell cocultures

0.4 μm pore polyester membrane transwells (Corning) were used for transwell cocultures. 2,000 A549 cells were seeded on the upper chamber and the 8000 GFP^+^ TS-KO cells were seeded on the lower chamber. The growth of TS-KO cells was monitored by Incucyte S3 to obtain the proliferation kinetics.

### Drug treatments

Carbenoxolone (CBX, Sigma) and SMIFH2 (Tocris) were first titrated in cells to determine the non-toxic concentrations. For cocultures, 60,000 parental cells and TS-KO cells were seeded per well in a 12 well plate in the determined concentration of the drugs (100 μM CBX and 50 μM SMIFH2), along with the respective vehicle controls (0.1 % NaCl for CBX, 0.1 %DMSO for SMIFH2). One day after the treatment, cells were trypsinized, counted and coculture was set-up as described in previous section. During the coculture cells were maintained in treatment conditions and proliferation was monitored using Incucyte real-time proliferation assay.

### Spheroid coculture

2,000 TS-KO cells per well were seeded in the presence or absence of parental cells treated with 100 μM CBX or 0.1 % NaCl in a 96-well ultra-low attachment plate. The plate was centrifuged at 300 rpm for 10 minutes at 20°C and then incubated at 37°C for three days until the spheroids attained a desired diameter of 200–500 μm. The spheroids were then re-treated with CBX (100 μM) or with 0.1% NaCl. Next, the plate was pre-cooled on ice, Matrigel matrix (Corning) was added to the spheroids, and the plate was centrifuged at 300 rpm for 5 minutes at 4°C. Then, the matrigel was polymerized in an incubator at 37°C for 30 minutes. The brightfield and green area of the spheroids were measured using a real-time spheroid invasion assay every 4 hours in Incucyte S3.

### Western blot

For western blots, 5–20 μg of protein were resolved on 12% bis-acrylamide gel and transferred on 0.45 μm PVDF membrane (Thermo). Membranes were blocked in 5% milk (Biorad) or 3% BSA (Sigma), both prepared in 1X TBS-T (1% Tween20 in Tris Buffer Saline). Membranes were then incubated with primary antibody (diluted in the respective blocking buffer) overnight at 4°C. Post incubation, membranes were washed with 1X TBS-T and incubated with HRP-conjugated secondary antibody for an hour at room temperature. After washing again with 1X TBS-T, ECL substrate (Thermo) was applied to the membrane and the bands were recorded on X-RAY films (Thermo) and developed and fixed manually. A list of antibodies and dilutions is provided in Extended Table 2.

### β-Galactosidase staining

A Senescence β-Galactosidase Staining Kit (Cell Signaling Technology) was used. In brief, cells were seeded at low confluence in a 6- or 12-well plate. 24 hours post seeding, the medium was changed, and the cells fixed and stained following the instructions from the manufacturer. Stained cells were visualized and imaged using an Olympus BX53 microscope. Positive cells and total cells were counted using ImageJ and positively stained cells were normalized to the total number of cells.

### siRNA transfection

Reverse transfection was done with Lipofectamine RNAiMAX Transfection Reagent (Thermo). The transfection complex was prepared by adding 80 nM siRNA to 2 µl transfection reagent in 200 µl serum free DMEM and incubated for 15 minutes. After incubation, the transfection complex was added to the bottom of 12-well plates, and 60,000 suspended in 800 µl and added to the wells on top of the transfection complex. Cells were incubated at 37°C in 5% CO_2_. 24 hours post transfection, cells were used to establish cocultures. Knock down was confirmed in the lysates that were collected 48 hours post transfection.

### Immunofluorescence

For immunofluorescence, 100,000 cells/well were seeded in 6-well plates on coverslips (VWR). For coculture conditions, H2B-fluorescent tagged TS-KO cells were seeded with untagged parental cells. One day post seeding, cells were fixed with 4% formaldehyde (VWR) for 5 minutes followed by 10 minutes of permeabilization with 1X PBS-T (0.1% Triton-X100 in 1X PBS). Cells were blocked in 3% blocking buffer (3% BSA in PBS-T) and incubated in primary antibody (diluted in blocking buffer) overnight at 4°C. Then, cells were washed with 1X-PBS and incubated with secondary anti-rabbit antibody conjugated with Alexa-488 or Alexa (diluted in blocking buffer) for an hour. Cells were then mounted in DAPI containing mounting medium and visualized using an Olympus BX53 microscope.

### FRAP for measuring coupling

Monolayers in 4-well ibidi imaging slides were loaded with calcein red-orange (Invitrogen, F10319) for 10 minutes in HEPES RPMI (Thermo). Medium was then replaced by fresh medium to remove unloaded dye. Confluent monolayers were imaged using a Zeiss LSM 700 confocal microscope. A cell in the middle of a confluent cluster was selected for bleaching (high-power 555 nm) until fluorescence at 555 nm excitation and emission >590 nm decreased by 50%. The fluorescence signal was measured in the central cell, neighboring cells, and more remote cells, and normalized to the initial intensity. Recovery was fitted to a monoexponential to calculate calcein permeability (in μm/minutes) from the product of the geometric perimeter and the area of the central cell, divided by the time constant of fluorescence recovery. Permeability from FRAP recording was calculated as described previously (*10*).

### *In vitro* quantification of C^13^ thymidylate

For dTMP^13^ quantification in TS-KO cells, A549 parental cells (puromycin sensitive) were seeded at a density of 2 million cells in a 10-cm culture dish. After 24 hours, medium was replaced with medium supplemented with 400 µM 1-^13^C-Serine (Sigma). 24 hours post C^13^ enrichment, 1 x 10^6^ A549 H2B-GFP TS-KO cells were seeded with C13 enriched parental cells. Cell growth was monitored using Incucyte S3. When the growth of the TS-KO cells was restored, 10 μg/ml puromycin was added to the culture to eliminate the puromycin sensitive parental cells. Following clearing of the parental cells form the culture, TS-KO cells were washed five times with 1X PBS and then 1 ml of prechilled extraction solution was added (Methol:Isonitrile:Water; 2:2:1). Cells were incubated on dry ice for 20 minutes and then scraped and collected in 2 ml Eppendorf tubes. After further homogenization on dry ice via a bead beater (FastPrep-24; MP Biomedicals, CA, USA) at 6.0 m/s (3 x 30 seconds, 5 minutes pause time) using 1.0 mm zirconia/glass beads (Biospec Products), samples were centrifuged at 15,000 × g, 4°C for 10 minutes using a 5415R microcentrifuge (Eppendorf) and supernatants were transferred. The remaining pellet was reextracted by addition of 500 µL of a mixture of H_2_O:MeOH:ACN (1:2:2, v/v) and bead beating (see conditions described above). After centrifugation, supernatants of corresponding samples were combined and dried under a stream of nitrogen (Organomation Microvap). Dried samples were reconstituted in 50 µL H_2_O:MeOH:ACN (1:2:2, v/v), vortexed, centrifuged, and transferred to an analytical glass vials. The LC-MS/MS analysis was initiated within one hour after the completion of the sample preparation.

### LC-MS/MS analysis

LC-MS/MS analysis was performed on a Vanquish UHPLC system coupled to an Orbitrap Exploris 240 high-resolution mass spectrometer (Thermo) in positive ESI (electrospray ionization) mode. Chromatographic separation was carried out on an Atlantis Premier BEH Z-HILIC column (Waters; 2.1 mm x 100 mm, 1.7 µm) at a flow rate of 0.25 mL/minutes. The mobile phase consisted of water:acetonitrile (9:1, v/v; mobile phase phase A) and acetonitrile:water (9:1, v/v; mobile phase B), which were modified with a total buffer concentration of 10 mM ammonium acetate. The aqueous portion of each mobile phase was adjusted to pH 9.0 via addition of acetic acid. The following gradient (20 minutes total run time including re-equilibration) was applied (time [minutes]/%B): 0/95, 2/95, 14.5/60, 16/60, 16.5/95, 20/95. Column temperature was maintained at 40°C, the autosampler was set to 4°C, and the sample injection volume was set to 5 µL. dTMP isotopologues were recorded via a targeted selected ion monitoring (tSIM) scan with a mass resolving power of 120,000 over a 5 Da isolation window set to m/z = 322.5 (scan time: 150 ms, RF lens: 70%). Ion source parameters were set to the following values: spray voltage: - 3500 V, sheath gas: 30 psi, auxiliary gas: 5 psi, sweep gas: 0 psi, ion transfer tube temperature: 350°C, vaporizer temperature: 300°C.

All experimental samples were measured in a randomized manner. Pooled quality control (QC) samples were prepared by mixing equal aliquots from each processed sample. Multiple QCs were injected at the beginning of the analysis to equilibrate the analytical system. A QC sample was analyzed after every 5th experimental sample to monitor instrument performance throughout the sequence. For determination of background signals and subsequent background subtraction, an additional processed blank sample was recorded. Data was processed using TraceFinder 5.1 and raw peak area data was exported for relative metabolite quantification.

### Cell-line derived xenografts

For cell line derived xenografts, 500,000 cells were suspended in 40 μl Matrigel (Corning)/1X PBS (1:1). Cells were subcutaneously injected in the flanks of 8–12 week-old female NSG (NOD scid gamma, Charles Rivers Laboratories) mice. Once palpable tumors were observed, mice were sacrificed, by cervical dislocation and tumors were excised and weighed. Tumor volumes were calculated from length and breadth of tumors obtained from caliper measurements using the formula π/6 x length^2^ x width. Mouse experiments were approved by The Danish Animal Experiments Inspectorate at Dyreforsoegstilsynet (License Number 2020-15-0201-00691) and all ethical considerations were followed. Number of mice have been specified for each experiment in their respective figure legends.

### CRISPR/Cas9-mediated lung tumorigenesis

Ultra-purified recombinant adeno-associated viral particles carrying gRNA for *Kras*-*p53* (AAV-KP) and *Kras*-*p53*-*Tyms*-*Tk1* (KPTT) were produced and obtained from VectorBuilder Inc., USA. For experiments B6J.129(B6N)-Gt(ROSA)*^26Sortm1(CAG-cas9*,-EGFP)Fezh/J^* mice were used which expressed Cas9 controlled by lox-Cre-lox (LSL) inducible site. Mice (8–40 weeks) were randomized for age and gender and a viral titer of 3×10^11^ was delivered via an oropharyngeal route to the lungs of C57BL/6 mice with cre-controlled expression of Cas9 (B6J.129(B6N)-Gt(ROSA)*^26Sortm1(CAG-cas9*,-EGFP)Fezh/J^*), resulting in two mouse genotypes: B/6.Kras^G12D/G12D^p53^Δ/Δ^ (KP) and B/6.Kras^G12D/G12D^p53^Δ/Δ^Ts^Δ/Δ^Tk1^Δ/Δ^ (KPTT). Mice were weighed weekly and sacrificed when they reached the humane endpoint defined by asphyxiation, 20% drop in body weight and immobility. Lungs were then collected and fixed in 4% buffered formalin and stained for H&E. For CBX treatment, KP and KPTT mice were subjected to biweekly treatment of 40 mg/kg CBX or corresponding volume of 0.9% NaCl (vehicle control). Mouse experiments were approved by The Danish Animal Experiments Inspectorate at Dyreforsoegstilsynet. License Number 2020-15-0201-00607. All ethical consideration were followed.

### Flow cytometry and sorting

For FACS of A549 H2B-GFP TS-KO cells in coculture, cells were trypsinized and resuspended in PBS. GFP^+^ TS-KO cells were gated using a non-GFP parental control. Sorted TS-KO and parental cells were reseeded for cocultures and monitored in Incucyte S3 for proliferation. For counting GFP^+^ KPTT cells, cells were trypsinized, resuspended in PBS, and run on a FACS Aria-III flow cytometer (BD). GFP+ cells were gated on the FITC channel. Data was analyzed on FlowJo.

## Quantitative-image-based cytometry (QIBC)

For QIBC analysis a coculture was set up in 6 well plates on coverslips (12,000 TS-KO cells and 4000 parental cells, TS-KO cells were tagged either with H2B-GFP or H2B-RFP). Proliferation was monitored in Incucyte S3 and when growth was observed in TS-KO cells in coculture, cells were fixed with 4% formaldehyde buffered with 0.5–1.5% methanol (VWR, 9713) for 12 minutes at room temperature, washed with ice-cold 1X PBS and permeabilized with 0.1% Triton-X in PBS. For CldU treatment, cells were incubated with 10 µM of CldU for 48 hours before co-culture for another 2-3, fixed and permeabilized as outlined above. All antibodies were diluted in Dulbecco’s modified Eagle’s medium (DMEM, high glucose, Glutamax) containing 10 % FBS. Coverslips were then incubated with primary antibodies for 1 h in dark at room temperature and washed with three times each with 1X PBS and then Coverslips were washed with PBS containing 0.2 % Tween (Sigma-Aldrich). Primary antibodies with the dilutions used were: p21 (1:500 rabbit), p-TBK1 (1:500, rabbit), and γH2AX (1:2,000, mouse, Biolegend, 613402). Next, secondary antibody conjugates of Alexa Fluor 488 or 568 for mouse (A-11029 or A-11031) and rabbit (A-11034 or A-11036) (ThermoFisher Scientific) in 1:2,000 dilutions were used. Secondary-antibody incubations were performed at room temperature for 30 minutes and were supplemented with 4′,6-diamidino-2-phenylindole dihydrochloride (DAPI, 0.5 μg/ml) to counter-stain DNA. After three washes in PBS, coverslips were washed twice in distilled water, dried and mounted in 4.5 μl Mowiol-based mounting medium (containing Mowiol 488 (Calbiochem)/glycerol/Tris-HCL, pH 8.5). For QIBC, images were acquired with a ScanR inverted microscope high-content screening station (Olympus) equipped with wide-field optics, air objective, fast excitation and emission filter-wheel devices for DAPI, FITC, Cy3, and Cy5 wavelengths, an MT20 illumination system, and a digital monochrome Hamamatsu ORCA-Flash 4.0LT CCD camera. Images were acquired in an automated fashion with the ScanR acquisition software (Olympus, 3.4.1). In total, 81–144 images were acquired containing at least 1,000 cells per condition. Acquisition times for the different channels were adjusted for non-saturated conditions in 12-bit dynamic range, and identical settings were applied to all the samples within one experiment. Images were processed and analyzed with ScanR analysis software. First, a dynamic background correction was applied to all images. The DAPI signal was then used for the generation of an intensity-threshold-based mask to identify individual nuclei as main objects. This mask was then applied to analyze pixel intensities in different channels for each individual nucleus. For analysis of sub-nuclear foci, additional masks were generated by segmentation of the respective images into individual spots with intensity-based or spot-detector modules included in the software. Each focus was defined as a sub-object, and this mask was used to quantify pixel intensities in foci. After this segmentation of objects and sub-objects, the desired parameters for the different nuclei or foci were quantified, with single parameters (mean and total intensities, area, foci count, and foci intensities) as well as calculated parameters (sum of foci intensity per nucleus). These values were then exported and analyzed with TIBCO Spotfire Software, version 12.2, to quantify absolute, median, and average values in cell populations and to generate all color-coded scatter plots. Within one experiment, similar cell numbers were compared for the different conditions (at least 1000 cells), and for visualization low x-axis jittering was applied (random displacement of objects along the x axis) to make overlapping markers visible.

### RNA-sequencing

Total RNA was isolated as per the manufacturer’s protocol (Qiazol) from A549 parental cells, A549 TS-KO and coculture conditions. Similarly, total RNA was isolated from lung tumor tissues of KP and KPTT mice (tumors were stored in RNase later and processed simultaneously). 500 ng of total RNA was diluted in 25 μl of nuclease-free water to construct the sequencing libraries using a NEBNext Ultra RNA Library Prep Kit for Illumina followed by paired-end sequencing using a NovaSeq 6000 platform (Illumina), as described by the manufacturer’s protocol. Fastq files generated from the sequencing were subjected to quality control analysis using the FastQC package. Alignment of the reads to the human reference genome (GRCh38 release 105) along with the respective human gene annotation file was performed using STAR aligner v2.7.9a. The GRCm39 release 108 was used as the reference genome for the mouse genome alignment. After the alignment, the reads were counted using the Feature counts function in the Rsubread v2.10.5 package in R 4.2.1 and subjected to differential gene expression analysis using the DESeq2 v1.36.0 package in R with an adjusted p-value<0.001 and a fold change of 2.

### Single-cell RNA-sequencing

Single-cell RNA-sequencing data of tumors from lung cancer patients were obtained from the public repository Gene Expression Omnibus (GSE148071, N = 42). The analysis was performed as described previously (*18*). In brief, the obtained raw data was subjected for quality control analysis and cells with a number of genes expressed < 200 & > 5000, UMIs with > 30,000 number of molecules, and > 30% of mitochondrial content were removed. The filtering of cells was performed for each patient’s sample and also in a pooled manner. Two patient samples (patient 19 and patient 27) were excluded from the analysis as the raw data contained duplicate gene names. Seurat v4.2.1 was used to normalize the counts for the library size and identified 600 variable genes to scale the data for the principal component analysis (PCA). Cell clusters were identified with the first 20 principal components from PCA and using a Lovain algorithm with a resolution set to 1.0. The identified cell clusters were visualized using uniform manifold approximation and the projection (UMAP) dimensionality reduction method. Cell clusters were scored and assigned to one of 11 major cell types based on the manual inspection of the cell type markers’ normalized expression level in UMAP, and also with the differentially expressed genes level using the FindAllMarkers function in Seurat. The highest scored cell type was assigned to each cluster. Markers used for the cell type assignment were: Endothelial cells (*CLDN5*, *VWF* and *PECAM1*), fibroblasts (*COL1A1*, *COL1A2* and *DCN*), neutrophils (*CSF3R*, *S100A8* and *S100A9*), mast cells (*GATA2*, *TPSAB1* and *TPSB2*), alveolar cells (*CLDN18*, *AQP4* and *FOLR1*), T cells (*CD2*, *CD3D*, *CD3E* and *CD3G*), epithelial cells (*CAPS* and *SNTN*), myeloid cells (*CD14* and *LYZ*), B cells (*CD79A* and *CD79B*), and follicular dendritic cells (*FDCSP*).

Cancer cell clusters were assigned as *EPCAM*-positive cells that was also negative for normal lung epithelial makers. In four patient samples, we did not find cancer cell clusters (patient 7, patient 11, patient 37 and patient 42). In the subsequent analysis, cell clusters assigned with the same cell type were pooled together. The number of cells in each cell type per patient is provided in the Supplementary Data S1. Next, cancer cell clusters were filtered to categorize each cancer cell based on *TYMS* and *TK1* expression levels. In brief, cells were assigned as follows: normalized expressions of *TYMS* > 0 and *TK1* = 0 as *TYMS^+^TK1^-^*, *TYMS* = 0 and *TK1* > 0 as *TYMS^-^TK1^+^*, *TYMS* > 0 and *TK1* > 0 as *TYMS^+^TK1^+^*, and finally *TYMS* = 0 and *TK1* = 0 as *TYMS^-^TK1^-^*. The number of cells in each of the categorized cell sub-types based on *TYMS* and *TK1* expression for each patient is provided in Supplementary Data S2.

With the assigned *TYMS* and *TK1* expression status for the cancer clusters, gene set enrichment analysis (GSEA) was performed utilizing the hallmark gene sets and other gene sets related to senescence and pyrimidine metabolism collected from MSigDB v2023.2.Hs and gene sets for chromosomal instability and immune response from previous reports (*44-46*). The details of the gene sets are provided in the Supplementary Data S3. A ranked gene list based on a log2 fold change was generated using FindMarkers function by comparing the *TYMS^-^TK1^-^*to the rest of the cancer cell types (*TYMS^+^TK1^-^*, *TYMS^-^TK1^+^* and *TYMS^+^TK1^+^*). GSEA was performed using fgsea package in R for the rank ordered gene list with the obtained gene sets in each patient sample. Significant gene sets were filtered with a p-value < 0.05 and the number of occurrences of each gene set across patient samples (N = 36) were tabulated. With a threshold of more than 10 patient samples, the gene sets were considered to be either enriched or diminished in *TYMS^-^TK1^-^* cells compared to the other cell states (*TYMS^+^TK1^-^*, *TYMS^-^TK1^+^* and *TYMS^+^TK1^+^*). A detailed list of enriched gene sets in each patient sample is provided in Supplementary Data S4. In case of proliferative marker analysis, cancer cells positive for *MKI67* marker (*MKI67* > 0) were filtered and analyzed for the proportion of different cell states based on *TYMS* and *TK1* expression levels. The number of each cell subtype status in the *MKI67* subset population is listed in Supplementary Data S5.

### Autodocking of GJA1 with dTMP

The structures of the Cx43 receptor in a fully closed state with the N-terminal gate closed (GCN) and a fully open state with the N-terminal lining the pore (PLN) were obtained from the Protein Data Bank with PDB IDs 7F94 and 7XQJ, respectively^52^. Missing loops were built by homology modeling using the Swiss Model server^53^. The ligand, thymidine-5’-phosphate, was obtained from the PubChem database^54^. Its protonation state was calculated using MarvinSketch. Both the ligand and receptor structures were imported into the Molegro Virtual Docker suite for molecular docking studies^55^. For the ligand, charges were assigned according to its protonation state, and all rotatable bonds were set as free. In both GCN and PLN conformations, blind docking was initially performed, followed by target docking. The Simplex Evolution algorithm and MolDock Score were used as the search algorithm and scoring function, respectively. Finally, the results were analyzed using the PLIP server^56^, and intermolecular interactions were visualized using PyMol^57^.

### Neighborhood analysis of Visium spatial sequencing

Spatial data was downloaded from X10 genomics (https://www.10xgenomics.com/datasets/human-lung-cancer-ffpe-2-standard). Sample id CytAssist_FFPE_Human_Lung_Squamous_Cell_Carcinoma. Spatial coordinates for Visium spots were extracted from the Seurat object and each spot was annotated with their transcription phenotype as *TYMS^+^TK1^+^*, *TYMS^+^TK1^-^*, *TYMS^-^TK1^+^*, or *TYMS^-^TK1^-^* depending on the expression of TYMS and TK1 genes. For every spot, the phenotypes of neighboring spots within a specified radius were counted using a fixed radius nearest neighbor method using the R dbscan package (*47*). For each phenotype, the mean number of neighbors was calculated across all spots expressing that phenotype. The neighborhood analysis was repeated for increasing radii (from 1 to 10) where the number of neighbors surrounding a given spot is calculated as Σ6n where n is the radius. Note that only those spots with complete neighbors (i.e. the number of neighbors must be equal to Σ6n, which is not the case with spots closer to the slide boundaries) were used for calculating the mean number of neighbors by phenotype; this was subsequently plotted as a percentage of cells of each phenotype surrounding a cell of given phenotype. Analysis was done in R v4.4 a.

## Statistical analysis

All statistical analyses were performed using GraphPad Prism 10, unless stated otherwise. Comparison-groups are indicated either in the figures with lines or stated in the figure legends. P-values of statistical significance are illustrated as follows: *<0.05; **p<0.01; ***p<0.001, ns represents non-significant results. Each experiment is the representative of at least three independent repeats.

## Extended Figures

**Extended Fig. 1.**
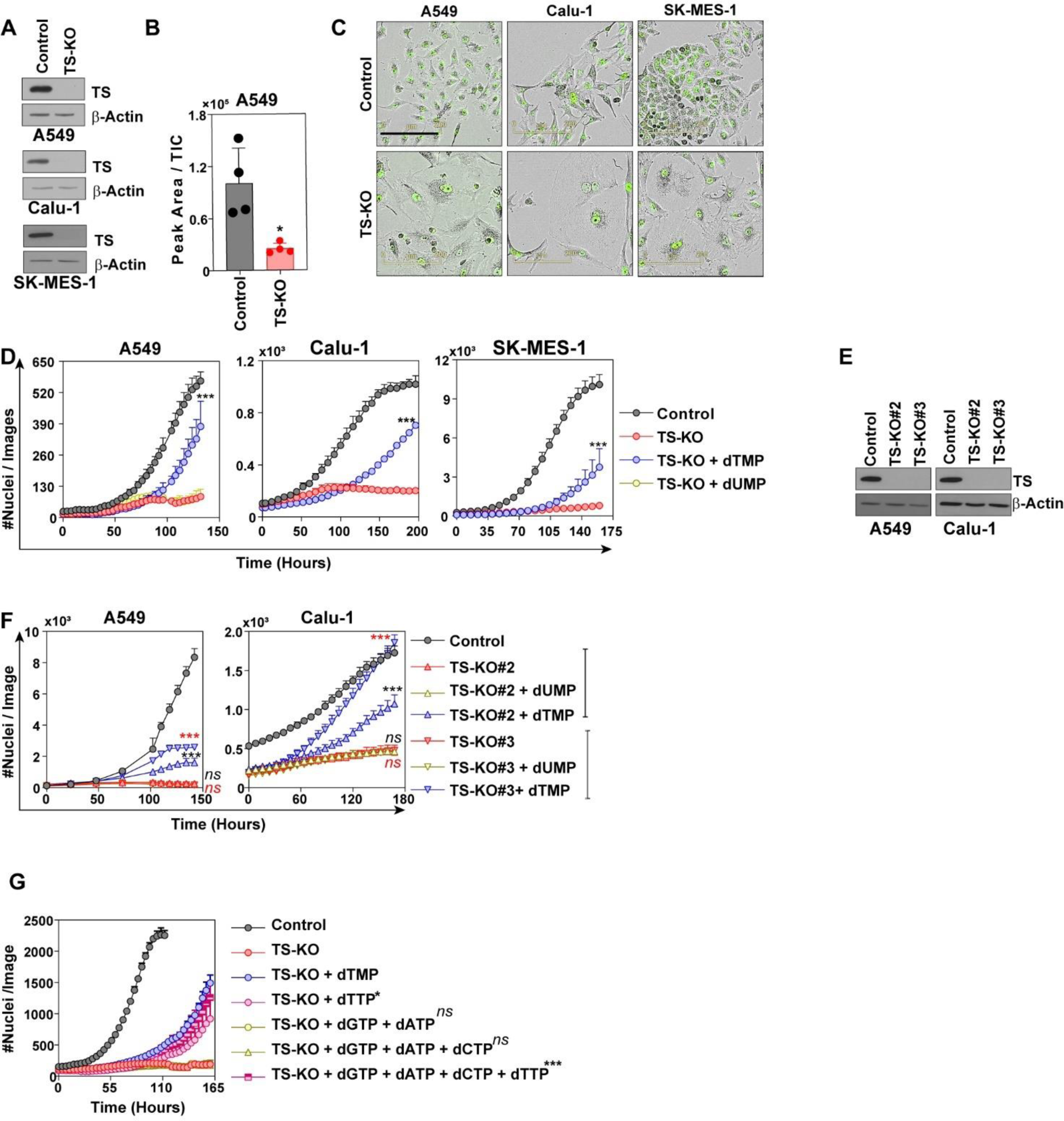
de novo deficient model dependent on exogenous dTMP.(A) Western blot quantification of TS-KO in indicated cell lines. **(B)** Mass-spec quantification of dTMP in A549 TS-KO cells compared to the control cells. Each bar represents mean ± SD of five replicates. Significance is calculated by Student’s t-Test. **(C)** Images from (B) showing TS-KO cells with enlarged morphology. Scale bar represents 200 μm. **(D)** Growth curve highlighting dependency of TS-KO cells on 10 μM dTMP supplementation in comparison to 10 μM dUMP (only A549). Significance is compared to TS-KO cells. Each point represents mean ± SD of three replicates. Significance is Two-Way ANOVA and Tukey’s Multiple Test. **(E)** KO of TS with non-overlapping gRNAs for *TYMS* other than what has been shown in Fig 1C and Extended Fig. 1A. **(F)** Treatment of the cells in (E) with 10 μM dTMP and 10 μM dUMP. Each point represents mean ± SD of three replicates. Significance compared to TS-KO calculated by Two-way ANOVA and Tukey’s Multiple Test. (**G**) Treatment of TS-KO cells with different combination of dNTPs, showing that only dTTP supplementation has positive effect on their growth. Each point represents mean ± SD of three replicates. Significance compared to TS-KO calculated by Two-way ANOVA and Tukey’s Multiple Test.

**Extended Fig. 2.**
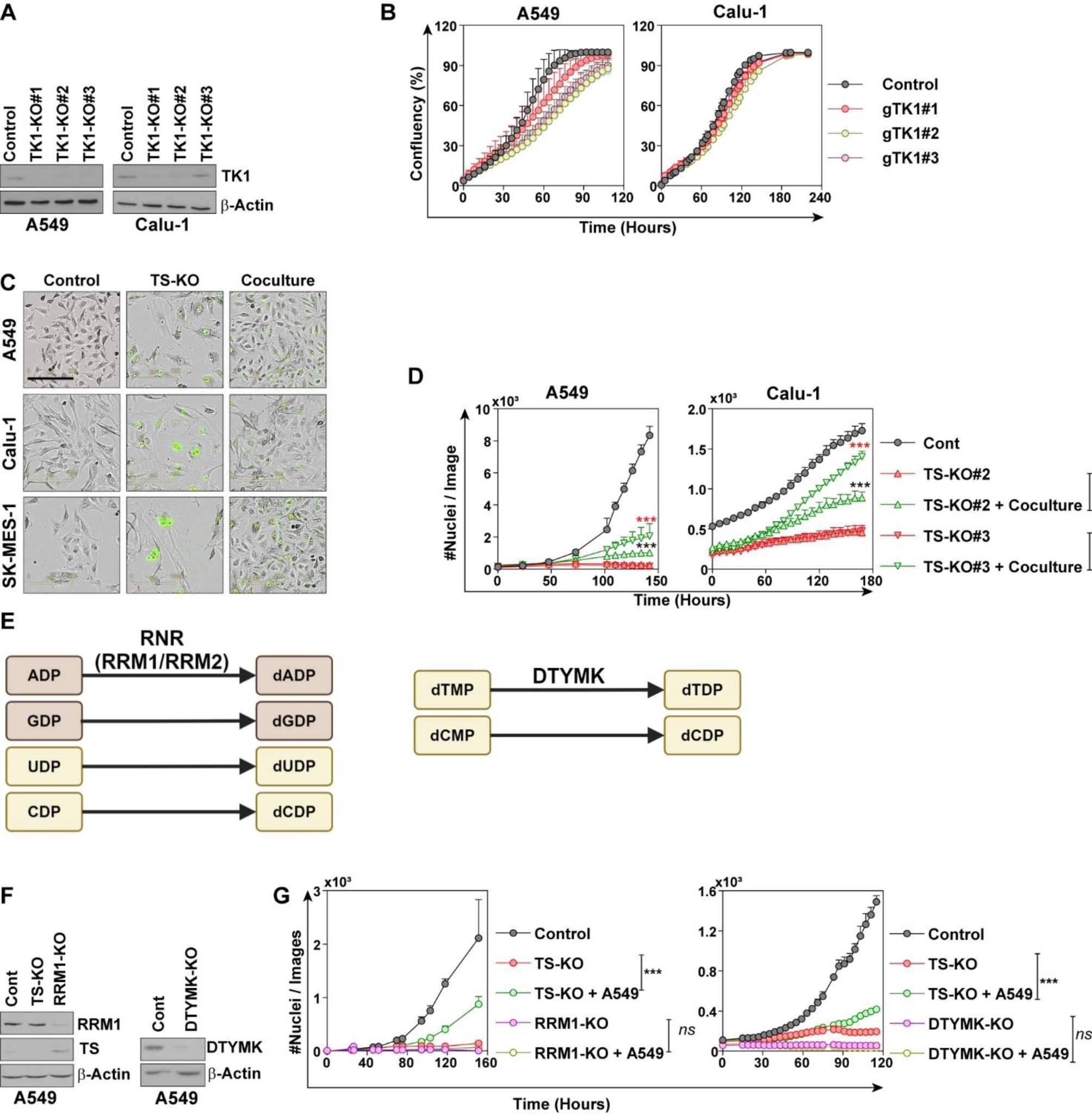
Collective proliferation is restricted to transfer of dTMP. **(A)** Western blot depicting CRISPR/Cas9 mediated knockout of *TK1* using multiple sequences in A549 and Calu-1 cells. **(B)** Proliferation curve showing growth in TK1-KO cells compared to the controls. Each point represents mean ± SD of three replicates. **(C)** Images from Fig. 1C showing that TS-KO cells regain normal morphology in coculture with parental cells. Scale bar represents 200 μm. **(D)** Rescue of the growth arrest of A549 and Calu-1 cells knocked out with two independent gRNAs, other than that shown in the Fig. 1C. Significance, compared to TS-KO, is Two-way ANOVA with Tukey’s multiple comparison. Each point represents mean ± SD of three replicates. **(E-F)** Western blot quantification of *RRM1* and *DTYMK* KO in A549 to deplete all dNTP and pyrimidine deoxyribonucleotides respectively. **(G)** Growth curve quantifying the proliferation of cells in coculture with parental A549 cells after *RRM1-* and *DTYMK1*-KOs. Each point represents mean ± SD of three replicates. Significance compared between cocultures and their respective controls by Two-way ANOVA with Tukey’s multiple comparison.

**Figure S3.**
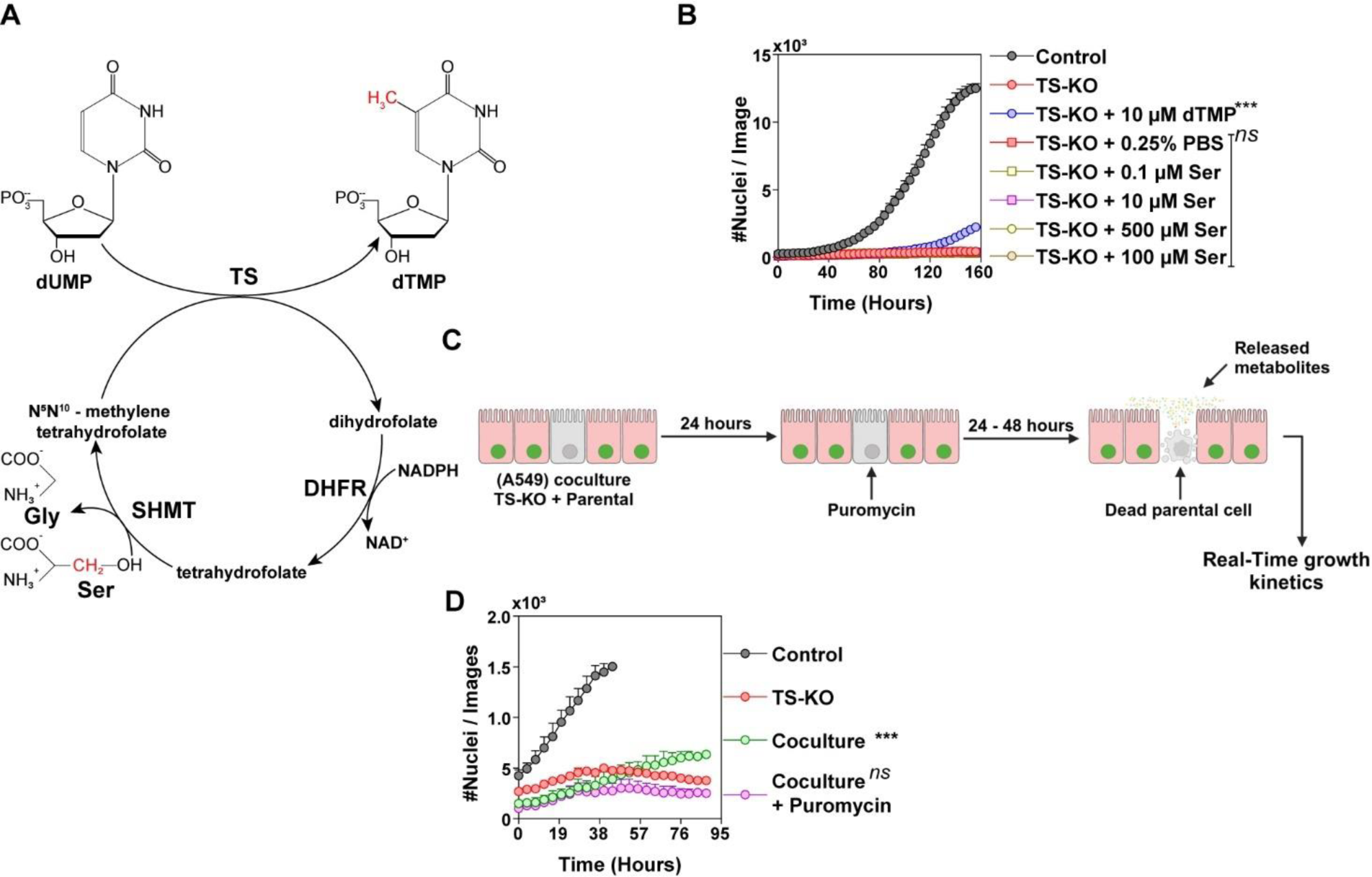
Serine as a source of dTMP detection. **(A)** TS-coupled folate cycle highlighting transfer of methyl group (red) from serine to dUMP for synthesis of dTMP. C13 labeled serine was used for detection of dTMP by mass spectroscopy shown in the Fig. 1F-G. **TS** = thymidylate synthase**, DHFR** = dihydrofolate reductase**, SHMT** = serine hydroxymethyl transferase. **(B)** Proliferation curve showing that serine did not directly contribute to support the growth of TS-KO cells. Significance is compared to TS-KO. Each point represents mean ± SD of three replicates. **(C)** Scheme depicting the proliferation kinetics of TS-KO cells in presence of parental cells treated with puromycin. **(D)** Proliferation curve showing that puromycin-killed parental cells did not contribute to the growth of TS-KO cells. Each point represents mean ± SD of three replicates. Significance is compared to TS-KO. Significance calculated in by Two-Way ANOVA with Tukey’s multiple comparison.

**Extended Fig. 4.**
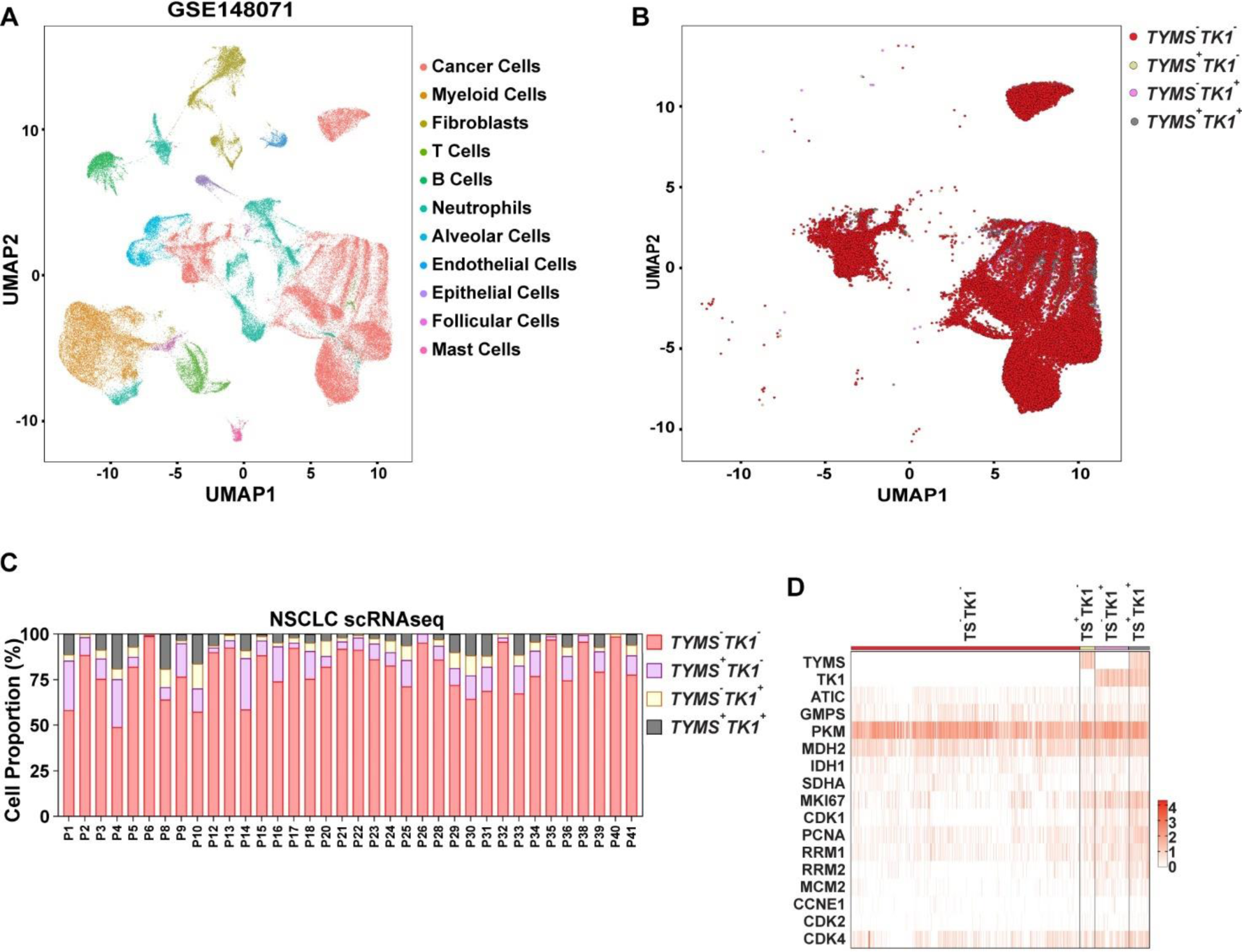
Quantification of *TYMS^-^TK1^-^*cells in scRNA seq DNA of NSCLC patients. **(A)** Single cells from 40 NSCLC patients clustered into different cell types (n = 40). Each dot represents a single cell. The number of cells is 88,672. **(B)** Representation of cells with both *de novo* (*TYMS^+^TK1^+^*), only *de novo* (*TYMS^+^TK1^-^*), only salvage (*TYMS^-^TK1^+^*) or neither (*TYMS^-^TK1^-^)* from cell types clustered from single cell data for 40 NSCLC patients. Each dot represents a cancer cells. The number of cancer cells is 39,787. **(C)** Patient-wise distribution of different population from (B). Each bar represents a single patient. **(D)** Expression of indicated metabolic genes and proliferation markers in TYMS^-^TK1^-^ cancer cells from 40 patients from (A) compared to other cell populations. *TYMS* = Thymidylate Synthase, *TK1*= Thymidine Kinase-1, Purine metabolism genes (*ATIC* = 5-Aminutesoimidazole-4-Carboxamide Ribonucleotide, *GMPS* = Guanine Monophosphate Synthase), glycolysis genes (*PKM* = Pyruvate Kinase M1/2), tricarboxylic acid cycle (MDH2 = Malate Dehydrogenase – 2, *IDH1* = Isocitrate Dehydrogenase (NADP(+))-1, *SDHA* = Succinate Dehydrogenase Complex Flavoprotein Subunit A), proliferation genes (*MKI67* = Marker Of Proliferation Ki-67, *CDK1* = Cyclin Dependent Kinase 1, *PCNA* = Proliferating Cell Nuclear Antigen, *RRM1* = Ribonucleotide Reductase Catalytic Subunit M1, *RRM2* = Ribonucleotide Reductase Catalytic Subunit M2, *MCM2* = Minichromosome Maintenance Complex Component 2, *CCNE1* = Cyclin E1, *CDK2* = Cyclin Dependent Kinase 2, *CDK4* = Cyclin Dependent Kinase 4).

**Extended Fig. 5.**
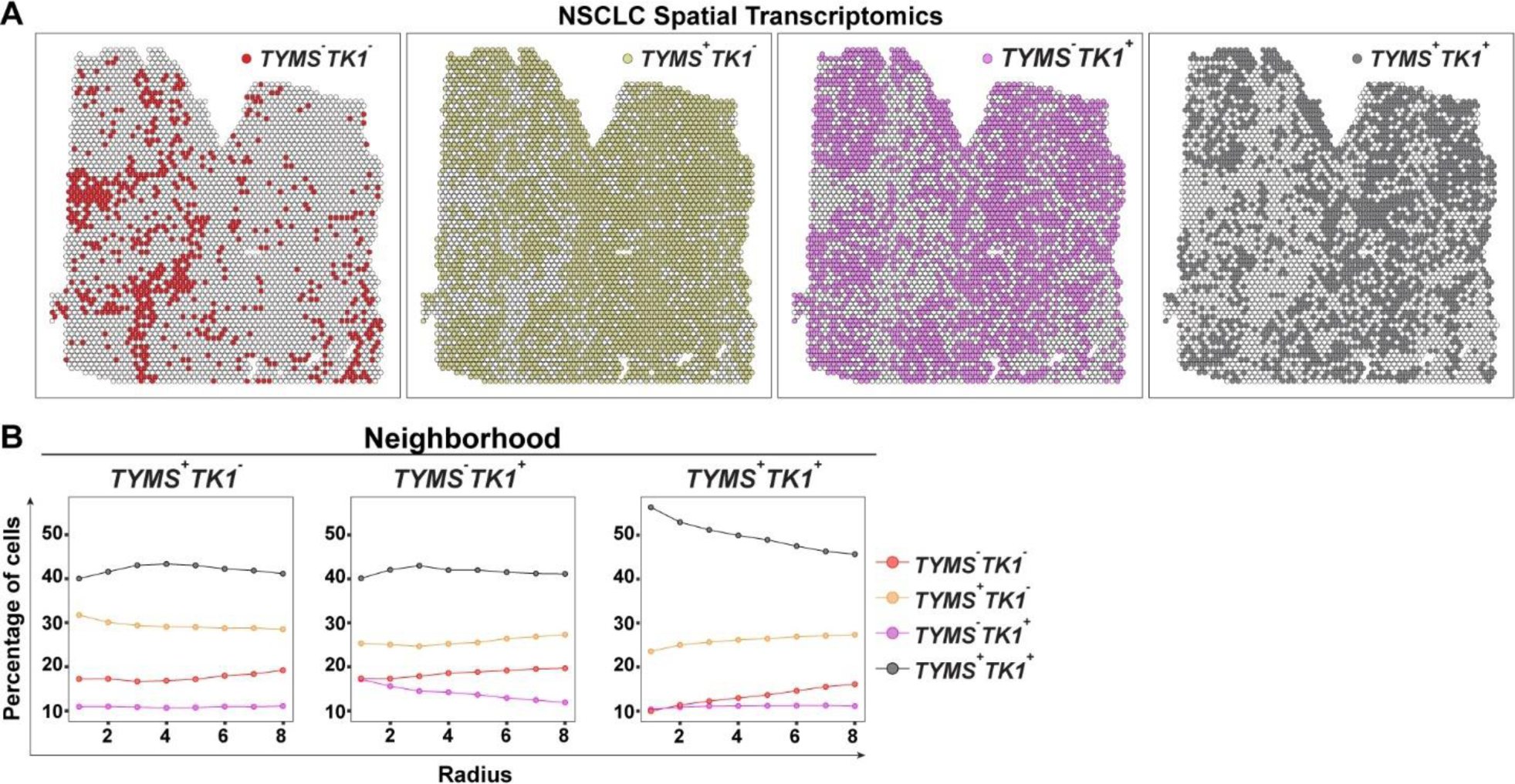
Spatial distribution of cells based on expression of *TYMS* and *TK1*. **(A)** Spatial distribution of indicated populations from Fig. 1I shown in individual map. **(B)** Neighborhood analysis of indicated population. See Fig. 1J.

**Extended Fig. 6.**
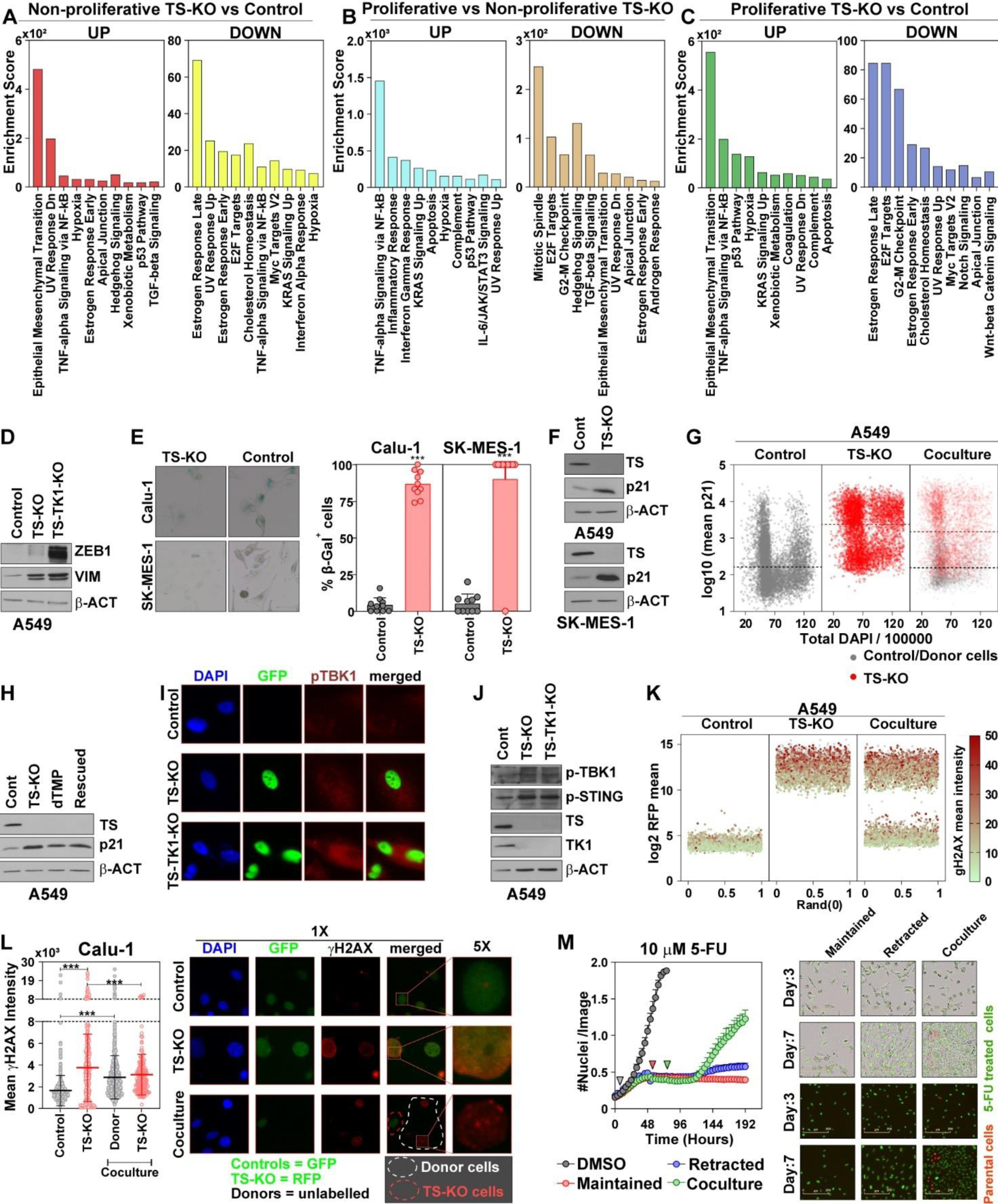
Pathophysiological features associated with collective proliferation. (A-C) Transcriptomic changes were globally analyzed in TS-KO cells and showing **A)** Differentially regulated pathways in TS-KO cells compared to control cells**, (B)** differentially regulated pathways in proliferative TS-KO cells (that were collected from coculture after puromycin induced death of parental donors) and non-proliferative TS-KO cells, and **(C)** differentially regulated pathways between proliferative TS-KO cells and control cells. **(D)** Western blot showing increased quantification of EMT makers in A549 cell line with dTMP deficiency. **(E)** Quantification of β-galactosidase, a marker of senescence in Calu-1 and SK-MES-1 cells. Bars represent mean ± SD of cells counted from 10 image fields. Statistical test is Student’s t-Test. **(F)** Increased expression of p21 in cells with TS-KO, as quantified by western blot. **(G)** Distribution of p21 in single cells from Figure 2C based on the total intensity of DAPI. Broken lines represent medians. **(H)** Quantification of p21 in A549 TS-KO cells in non-proliferative and proliferative (Rescued) states. **(I)** Representative images from Fig. 2F showing increased pTBK1 in cells with dTMP deficiency. **(J)** Quantification of cGAS-STING the effector proteins p-STING and p-TBK1 in TS-KO A549 cells. **(K)** Distribution of cells expressing γH2AX in TS-KO cells in coculture, showing increased mean γH2AX in RFP negative parental donor cells from Figure 2I. **(L)** Quantitation of γH2AX/cell in Calu-1 cells with TS-KO in coculture with donor parental cells. Dots represent cells, the longer line represents mean and shorter lines represent SD. Representative images, and 5X zoom in the inset show increased γH2AX in parental cells marked by white broken lines. Statistical analysis is Student’s t-Test. **(M)** Real-time proliferation of A549 cells treated with 10 µM 5-FU. Parental cells were added to the coculture after drug retraction showing that 5-FU induced growth arrest could be restored by coculture with parental cells. Grey arrowhead denotes the time of treatment, red arrowhead represents drug-retraction and green arrowhead indicates the time of coculture. Data is mean ± SD of three replicates. Significance is calculated by Two-Way ANOVA with Tukey’s multiple comparison.

**Extended Fig. 7.**
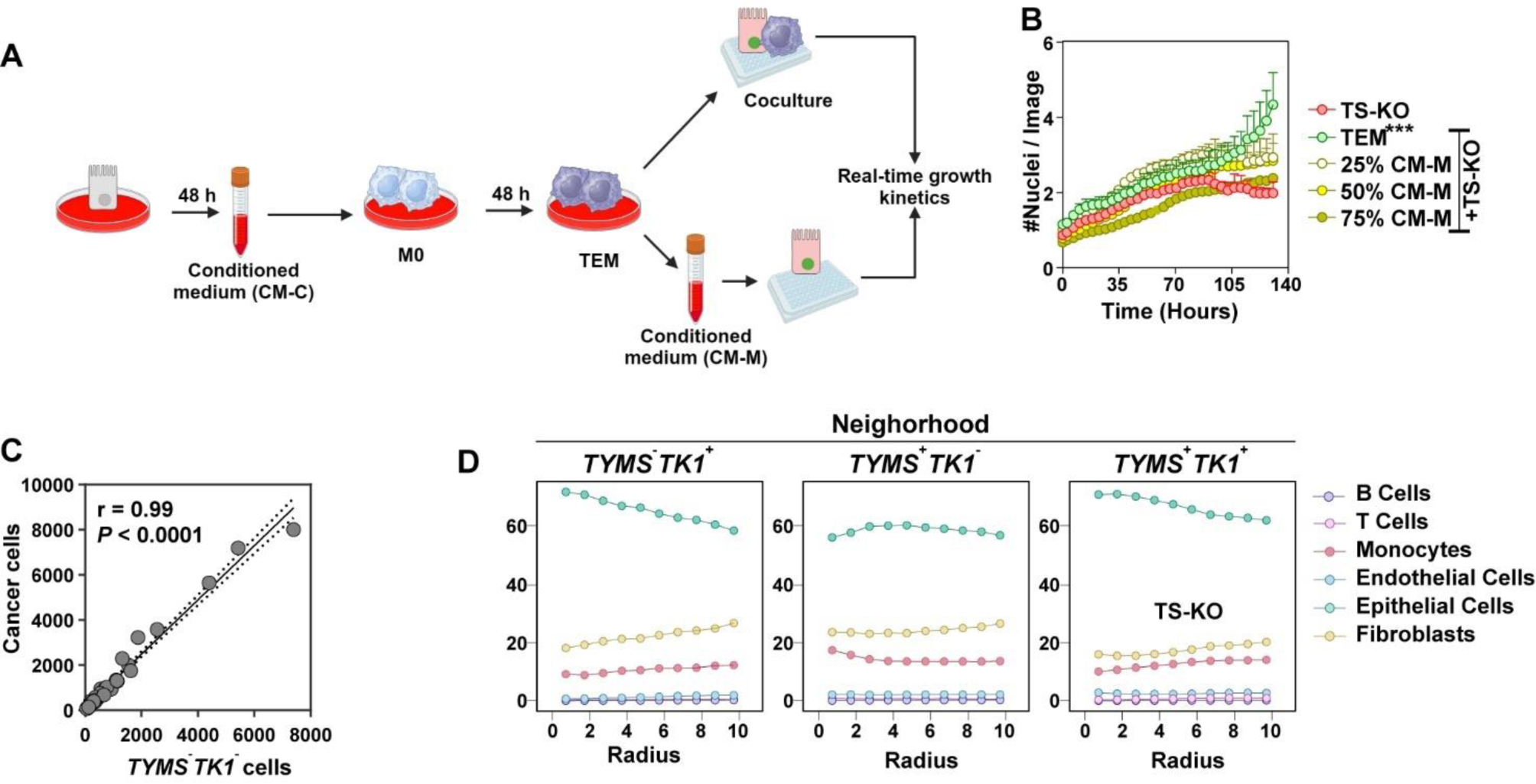
Contribution of tumor microenvironment (TME) cells to collective proliferation. **(A)** Schematic representation of the workflow followed for educating of M0 macrophages with cancer cells. Condition medium (CM-C) was collected from A549 cells after 48 hours of seeding. M0 macrophages were treated with CM-C for 48 hours and conditioned medium, referred to as CM-M, was then collected from tumor educated macrophages (TEMs). **(B)** Proliferation curve showing that CM-M in (A) has no effect on the growth of TS-KO cells. Data is mean ± SD of three replicates. Significance is calculated by Two-way ANOVA with Tukey’s multiple comparison. **(C)** Correlation between the numbers of total cancer cells and *TYMS^-^TK1^-^*from combined pool of cancer cells from patients in Extended Fig. 4. **(D)** Neighborhood analysis depicting proximity of *TYMS^-^TK1^-^* cells in Fig. 1I with different TME cells. See also Fig. 3M.

**Extended Fig. 8.**
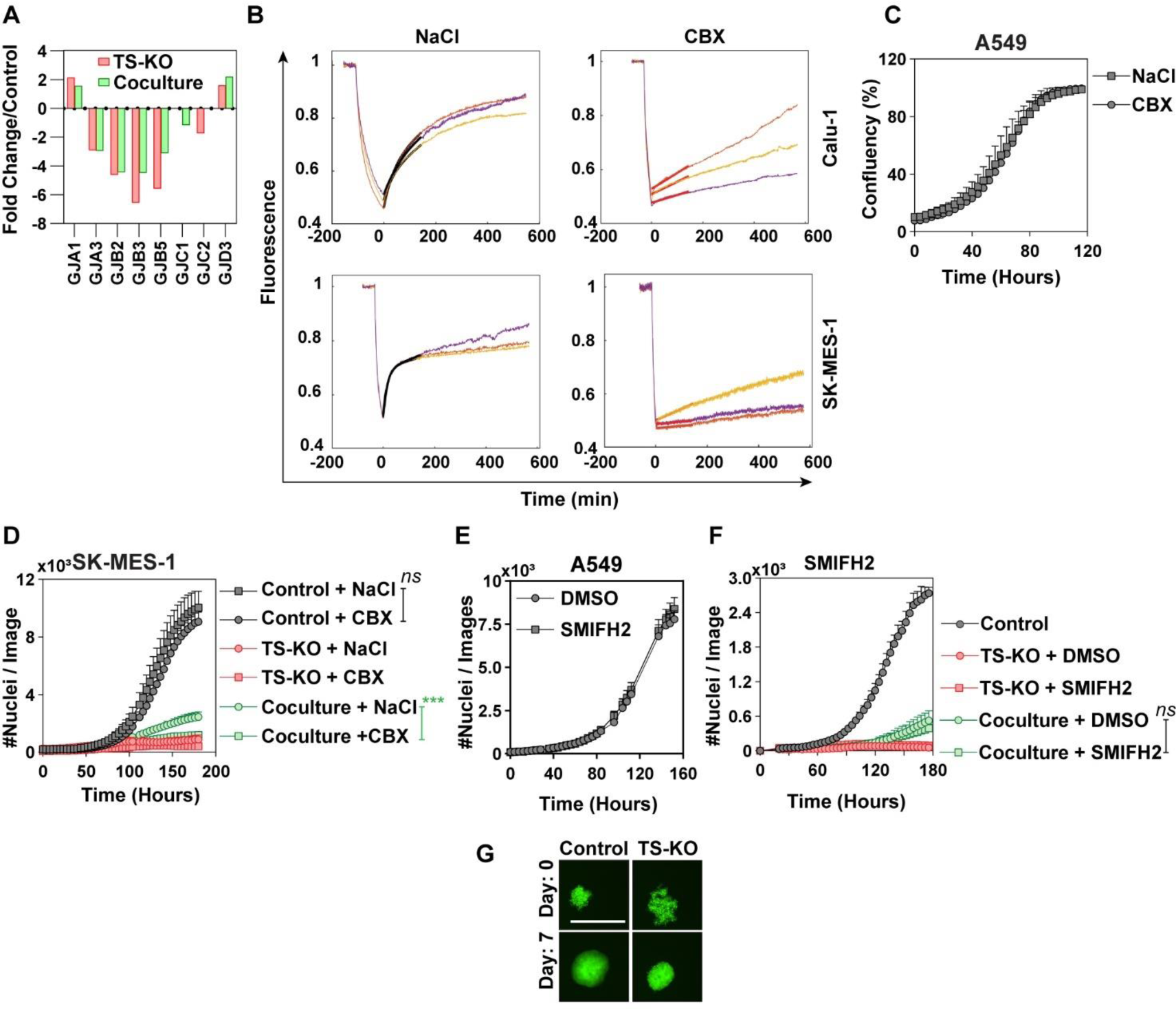
Gap junction connects cells for collective proliferation. **(A)** RNA sequencing analysis comparing TS-KO cells and puromycin selected TS-KO cells grown in coculture. GJA1 and GJD3 were two gap junction genes upregulated in both TS-KO and puromycin selected TS-KO cells. **(B)** Restoration of fluorescence in presence of either NaCl or CBX in Calu-1 and SK-MES-1 cell line from Fig 4A. **(C)** Proliferation curve showing that 100 μM CBX has no effect on the growth of A549 cells. **(D)** Collective proliferation reduced by 100 μM CBX in SK-MES-1 cells. Data is mean ± SD of three replicates. Significance is calculated by Two-Way ANOVA with Tukey’s multiple comparison. **(E)** Treatment of A549 cells with 50 μM of F-actin inhibitor SMIFH2, showing no effect on the growth of cells. **(F)** Collective proliferation of A549 cells with TS-KO was not affected in presence of 50 μM SMIFH2. **(G)** Representative images from Figure 4C, showing reduction of spheroid formation in A549 TS-KO cells. Scale bar represents 200 μm. All significances are calculated by Two-Way ANOVA with Tukey’s multiple comparison.

**Extended Figure 9.**
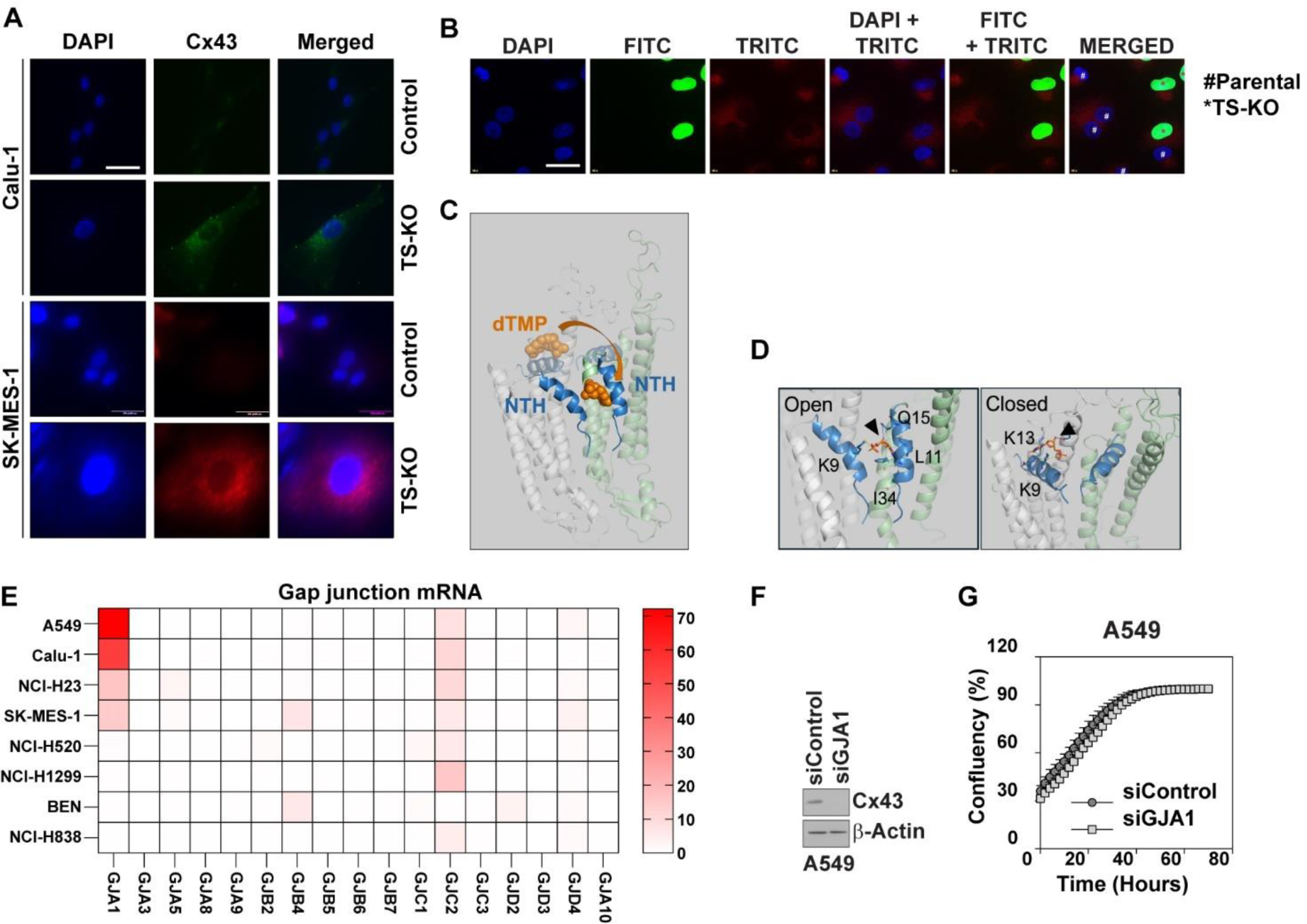
GJA1 is essential for collective proliferation. **(A)** Increased expression of Cx43 (*GJA1*) in in indicated cell lines quantified by immunofluorescence. Scale bar represents 200 μm. **(B)** Immunofluorescence staining of Cx43 in GFP^+^ TS-KO cells and non-GFP parental cells, showing increased expression of GJA1 in parental cells. Scale bar represents 200 μm. **(C-D)** Autodock based prediction of dTMP binding to the open conformation of GJA1. 9^th^ Lysine residue in Cx43 shows interaction with dTMP. Interactions have been indicated with black arrowheads. **(E)** Expression of GJA1 along with different gap junctions in selected NSCLC cell lines from CCLE data set. **(F)** Quantification of siRNA mediated knockdown of GJA1 in A549 cells. **(G)** Incucyte S3 quantification of proliferation in cells with GJA1 knockdown, showing that cells with siGJA1 in 4M has no proliferation defect. Each point represents mean ± SD of three replicates.

**Figure S10.**
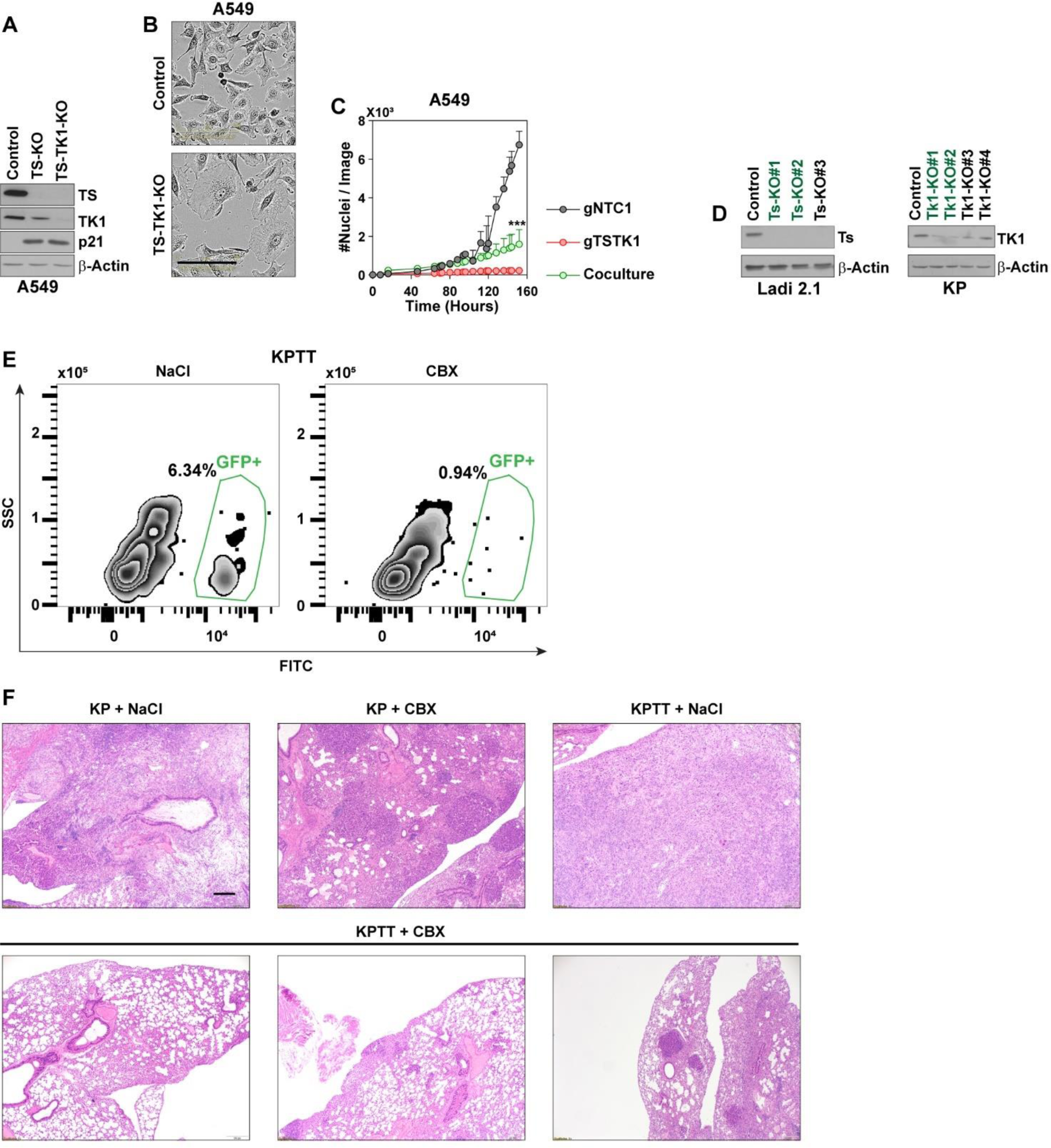
Growth of *TYMS^-^TK1*^-^ tumors are dependent on gap junctions. **(A)** Generation of A549 TS-TK1-KO cells used in Figure 5A showing loss of TS and TK1 along with increased p21. **(B)** A549 TS-TK1-KO cells showing senescence-like enlarged and flattened morphology. Scale bar represents 200 μm. **(C)** Real-time proliferation curve showing dependence on TS-TK1-KO cells on parental cells for growth. **(D)** Western blot showing quantification of Ts and Tk1 knockdown in cancer cells derived from C57BL/6 tumors. gRNAs #1 and #2, that are formatted in green, (both for Ts and Tk1) were used for generation of KPTT mice. (**E**) Flow cytometry quantification of GFP^+^ KPTT cells from coculture with Ladi 2.1 (a syngeneic lung cancer cell line that was established from KP on C57BL/6 background) cells in presence of either NaCl or 100 μM CBX. Quantification is shown in Fig. 5H. **(F)** Representative H&E images of lungs from Fig 5J, showing the tumor load in each group. Scale bar represents 100 μm.

---

**Extended Table 1.**
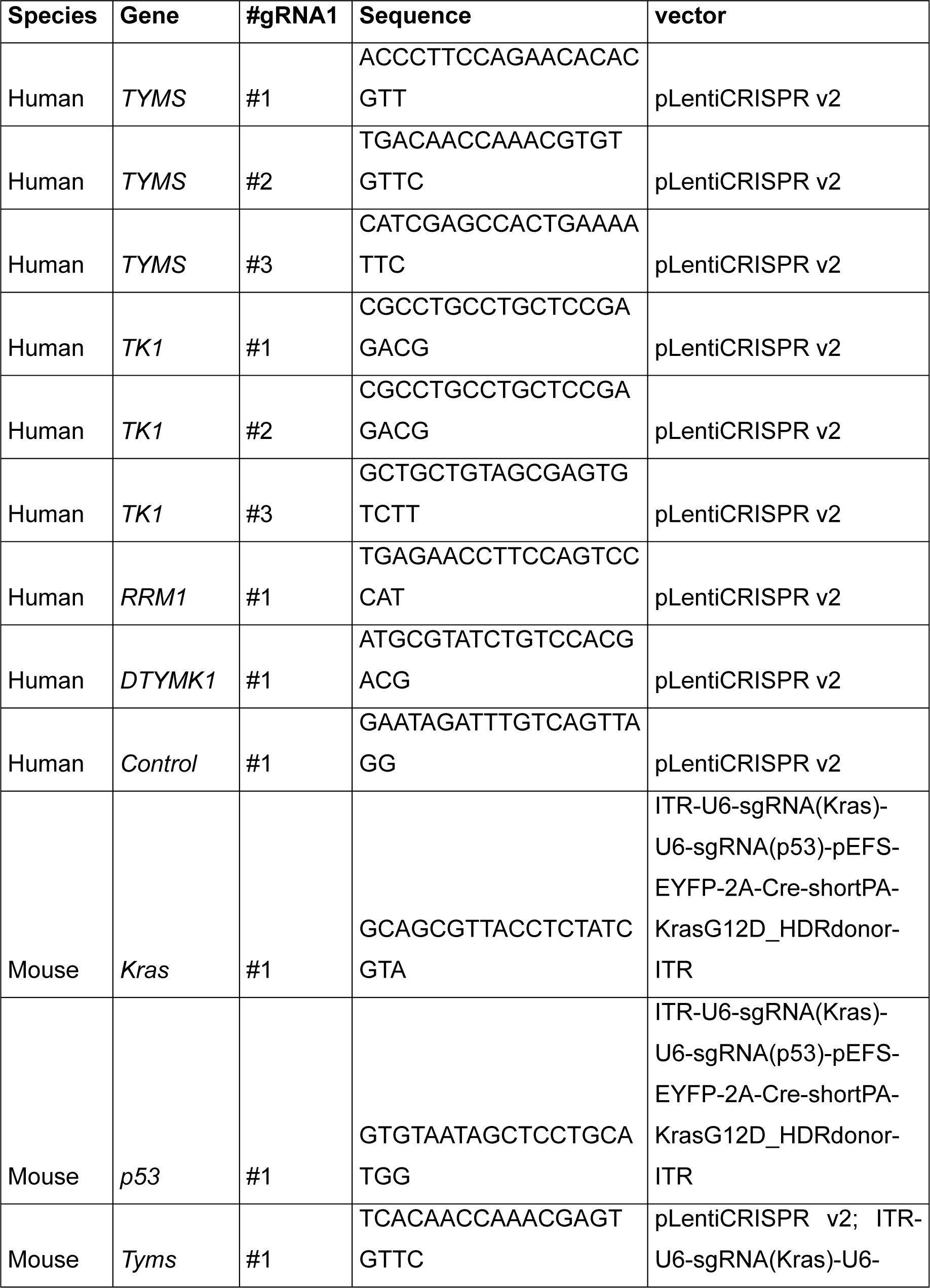
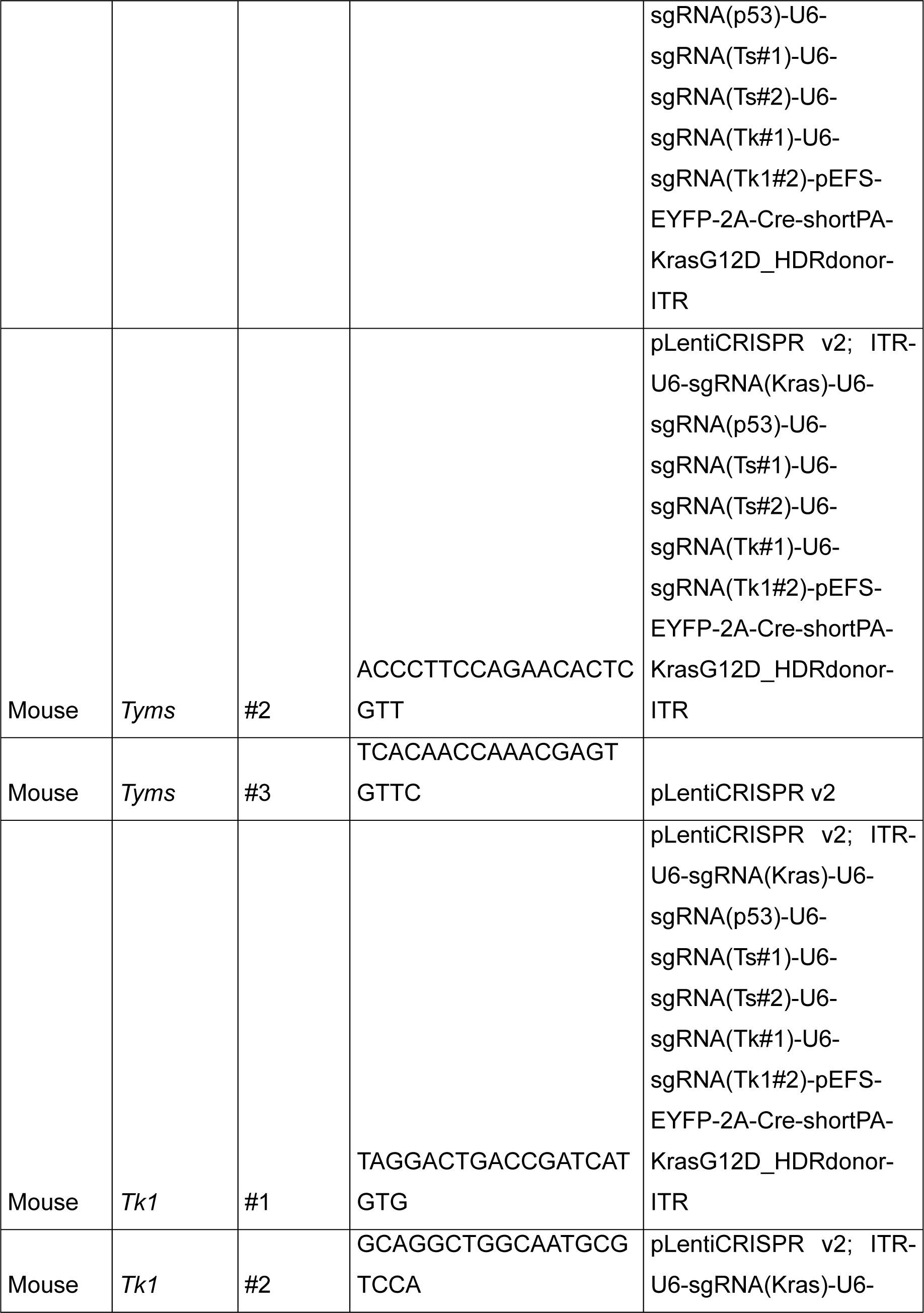
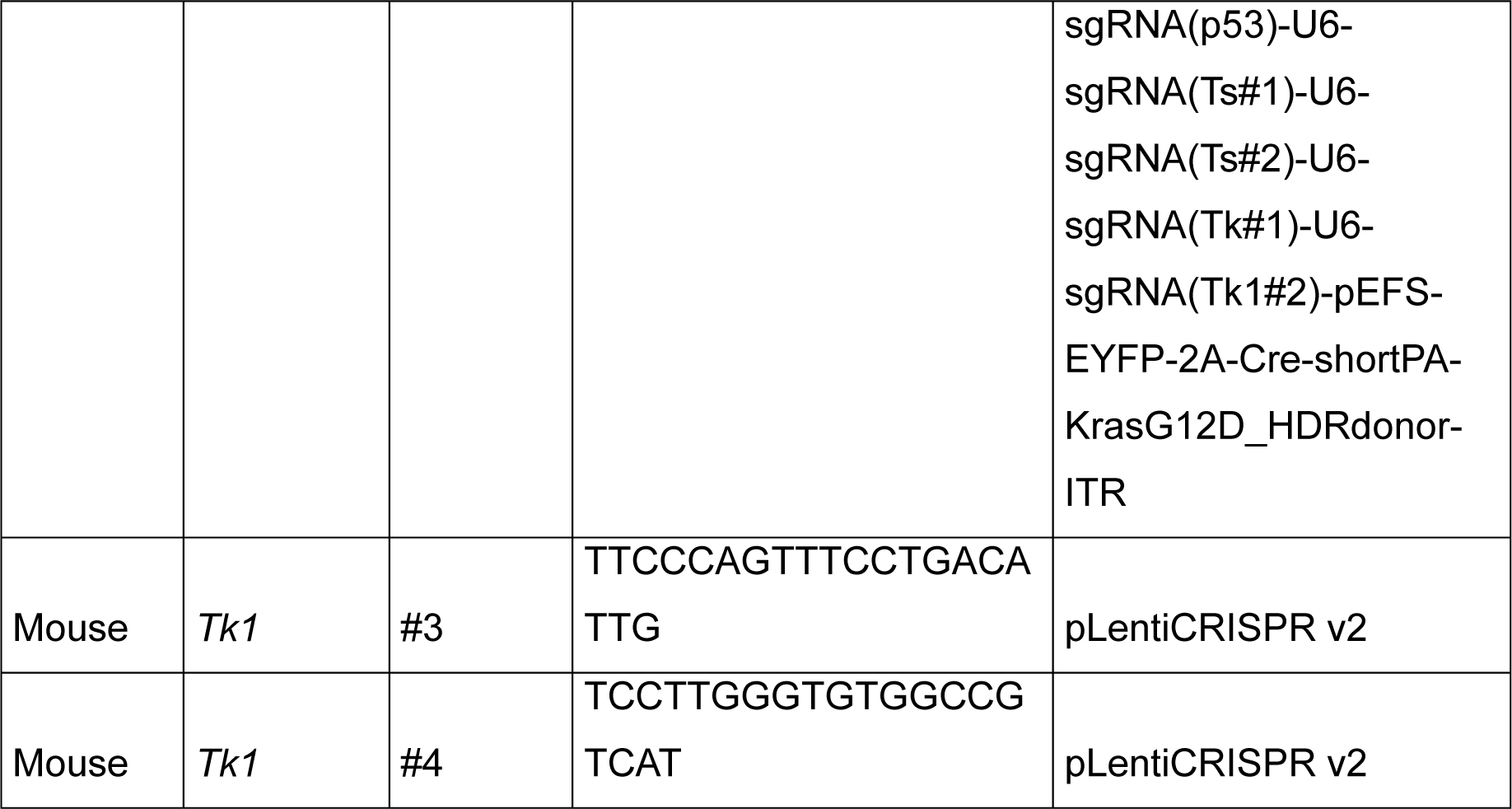
List of guideRNA Sequences.

**Extended Table 2.**
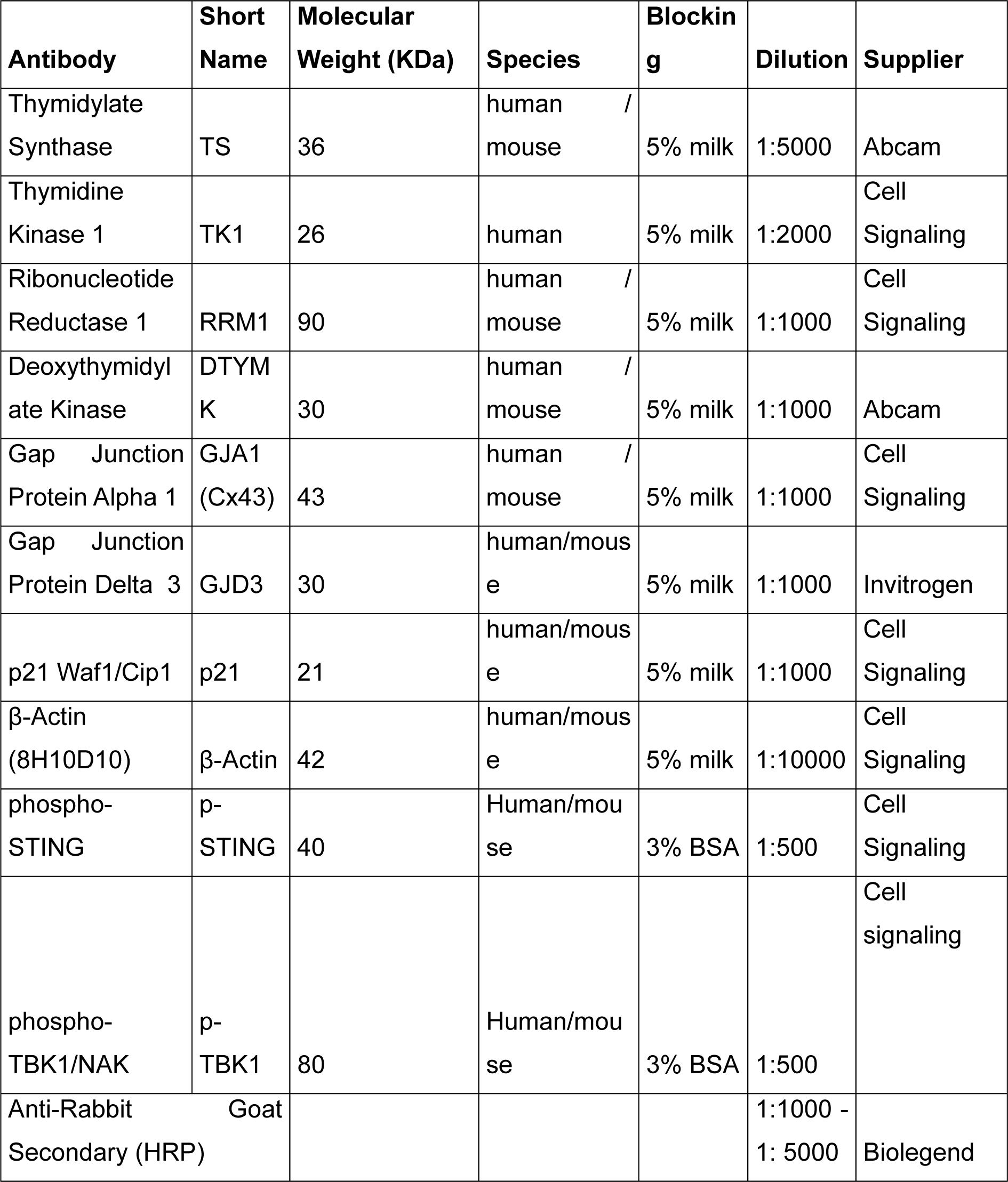
List of Antibodi.

